# The human microglia responsome: a resource to better understand microglia states in health and disease

**DOI:** 10.1101/2023.10.12.562067

**Authors:** Gijsje J.L.J. Snijders, Katia de Paiva Lopes, Marjolein A.M. Sneeboer, Benjamin Z. Muller, Frederieke A.J. Gigase, Ricardo A. Vialle, Roy Missall, Raphael Kubler, Towfique Raj, Jack Humphrey, Lot D. de Witte

## Abstract

Microglia, the immune cells of the brain, are increasingly implicated in neurodegenerative disorders through genetic studies. However, how genetic risk factors for these diseases are related to microglial gene expression, microglial function, and ultimately disease, is still largely unknown. Microglia change rapidly in response to alterations in their cellular environment, which is regulated through changes in transcriptional programs, which are as yet poorly understood. Here, we compared the effects of a set of inflammatory and restorative stimuli (lipopolysaccharide, interferon-gamma, resiquimod, tumor necrosis factor-alpha, adenosine triphosphate, dexamethasone, and interleukin-4) on human microglial cells from 67 different donors (N = 398 samples) at the gene and transcript level. We show that microglia from different anatomical brain regions show distinct responses to inflammatory stimuli. We observed a greater overlap between human stimulated microglia and human monocytes than with mouse microglia. We define specific microglial signatures across conditions which are highly relevant for a wide range of biological functions and complex human diseases. Finally, we used our stimulation signatures to interpret associations from Alzheimer’s disease (AD) genetic studies and microglia by integrating our inflammatory gene expression profiles with common genetic variants to map *cis*-expression QTLs (eQTLs). Together, we provide the most comprehensive transcriptomic database of the human microglia responsome.

**Highlights:**

- RNA-sequencing of 398 human microglial samples exposed to six different triggers.
- Microglia from different anatomical regions show distinct stimulation responses.
- Responses in human microglia show a greater overlap with human monocytes than murine microglia.
- Mapping of response Quantitative Trait Loci identifies interactions between genotype and effect of stimulation on gene expression.
- Our atlas provides a reference map for interpreting microglia signatures in health and disease.

## Introduction

In recent years there has been an increasing interest in understanding the dynamics of the microglial transcriptome (Gerrits et al. 2020; Sankowski et al. 2019). This interest has been boosted by genetic studies that have linked common and rare genetic risk factors for neurodegenerative diseases, such as Alzheimer’s disease (AD) and Parkinson’s disease (PD), to gene expression in microglia (Lopes et al. 2022; Kosoy et al. 2022; McQuade and Blurton-Jones 2019; Langston et al. 2022). Microglial changes have also been found in post-mortem brain tissue of patients with these diseases (Hopperton et al. 2018; Langston et al. 2022; Stefanova 2022). These include changes in density and morphology, as well as transcriptomic changes identified through bulk and single cell RNA-sequencing studies (Mathys et al. 2019; Hopperton et al. 2018; Olah et al. 2020; Gerrits et al. 2021; Sankowski et al. 2019; Sun et al. 2023). How these genetic associations and post-mortem changes should be interpreted in terms of microglia functions and the response of microglia to changes in their microenvironment, has proven to be challenging (Paolicelli et al. 2022).

As the primary resident immune cells of the brain, microglia perform a range of different functions (Matsudaira and Prinz 2022), including the production of inflammatory, restorative, or growth factors, phagocytosis, and more complex brain-specific functions through their interactions with neurons, including modulation of neurotransmission or synaptic pruning (Y. Liu et al. 2023). Like other myeloid immune cells, microglia express a range of receptors to detect changes in their microenvironment that require adaptation of their functions (Gosselin et al. 2017). If these receptors detect an alteration in their environment, microglia will rapidly respond with a change in morphology, migration, and modulation of their functions, which is regulated through the adaptation of transcriptional programs (Paolicelli et al. 2022).

Responses to microenvironmental changes have been extensively studied for rodent microglial cells (Lund et al. 2006; Pulido-Salgado et al. 2018; Das et al. 2018; Holtman et al. 2015) or for human monocyte and macrophage cultures (Fairfax et al. 2014; Gerrits et al. 2020; Alasoo et al. 2015; Xue et al. 2014; Cheng et al. 2019; De Jager et al. 2015), but are lacking for human microglia. The importance of understanding the transcriptional responses of human microglia is underscored by significant differences between microglia and other myeloid cells, as well as between microglia of different species (Butovsky et al. 2014; Geirsdottir et al. 2019; J. Melief et al. 2016). So far, only a few studies (Gosselin et al. 2017; Rock et al. 2005) investigated the effect of an altered microenvironment on human microglia (n = up to 19 donors). These studies found that the microglial transcriptome is highly dependent on its cellular and molecular context. Insights into the responsome of human microglial cells to inflammatory factors is necessary to further refine our understanding of context-dependent gene expression of microglial cells.

In this study, we describe the ***Microglia Genomic Atlas after STimulation (MiGASti)***, a genetic and transcriptomic data resource of 398 primary human microglia samples from 67 different human subjects (**Figure 1A)**. Microglia were either lysed directly after isolation or put in culture for 18 hours and exposed for 6 hours to a panel of inflammatory triggers. The response of microglia to these inflammatory and restorative stimuli was compared at the gene and transcript level, thereby defining specific microglial signatures across conditions. We compared our inflammatory transcriptional responses of microglia across different anatomical brain regions and to monocyte and murine mouse microglia. We further performed expression QTL analyses for cultured and stimulated microglial samples. Genetic variants that influence the inter-individual variation in responses to stimuli are called response eQTLs (reQTLs). Such genetic variants represent genetic effects that are modified by the environment via gene-by-environment interactions. Effects of a genetic variant on gene expression are context-specific and can be informative for disease. Finally, we compared our stimulation signatures to other published signatures to interpret genetic and transcriptomic data from Alzheimer’s disease as an example. To our knowledge, we are the first to investigate these large numbers of human microglial samples after exposure to different triggers and in relation to genetic effects.

**Figure 1.**
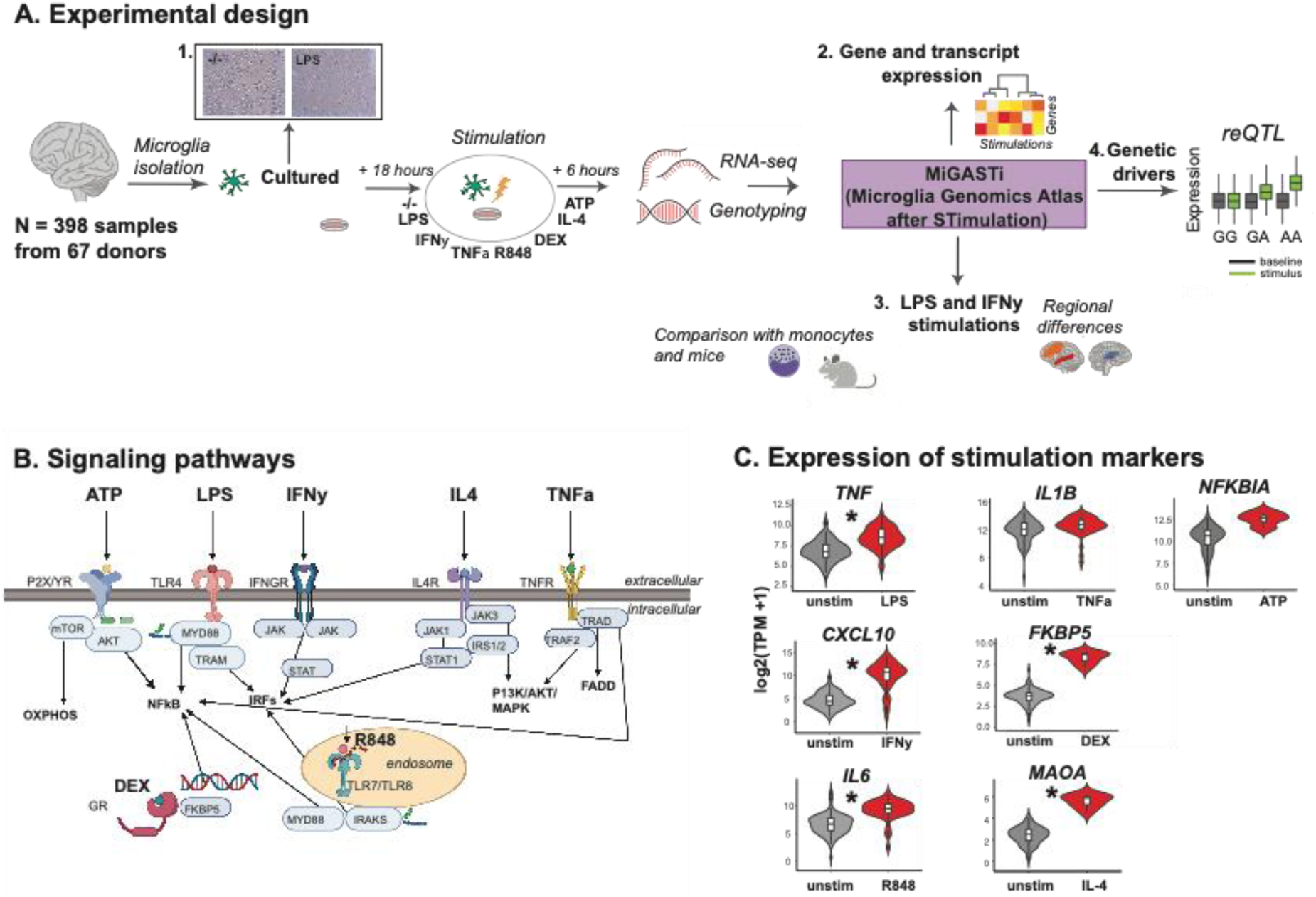
Overview of MiGASti. A) Primary human CD11b+ cells were isolated at autopsy from 67 different donors with neurological or psychiatric disease, as well as unaffected controls, generating a total of 398 samples from 7 brain regions. Samples were cultured for 24 hours to investigate the response of the cells to an altered inflammatory context, 1) the cells were exposed to lipopolysaccharide (LPS), imiquimod (R848), Interferon-gamma (IFNγ), tumor necrosis factor alpha (TNFα), Interleukin-4 (IL-4), Dexamethasone (DEX), or adenotriphosphate (ATP) for 6 hours prior to harvesting the cells. RNA was isolated and sequenced. Genome wide genotyping was performed using DNA isolated from all donors. The following analysis were performed with the MiGASti dataset: 2) gene and transcript related analysis with biological pathway analysis and functional enrichment. 3) Regional heterogeneity analysis for LPS and IFNγ and comparison with stimulated LPS and IFN© monocyte and mouse microglia data. 4) Response eQTL (reQTL) analysis. B) Schematic representation of the signaling cascades of the different environmental stimuli. C) Violin plots depicting expression in transcripts per million (TPM) for representative (anti)-inflammatory genes TNF, CXCL10, IL6, IL1B, FKBP5, MAOA, NFKBIA, that are known to be upregulated after stimulation with respectively LPS, IFNγ, R848, TNFα, DEX, IL-4 or ATP in the stimulated compared to the unstimulated samples. *Statistical significance based on adjusted P-values (FDR) derived from the differential expression results.

## Results

### A database of transcriptional responses of microglia to inflammatory and restorative triggers

We collected 483 human microglial samples from 87 different donors. Microglia were isolated from fresh post-mortem tissue using CD11b-beads. Microglial cells were directly lysed after isolation (*referred to as ‘uncultured’*), or they were cultured for 24 hours and then lysed (*referred to as ‘cultured’)*. To investigate the response of the cells to an altered inflammatory context, the cells were exposed to lipopolysacharriden (LPS), Interferon-gamma (IFNγ), Resiquimod (R848), Tumor-necrosis factor alpha (TNFα), adenotriphosphate (ATP), interleukin-4 (IL-4), or dexamethasone (DEX) or baseline (negative control) for 6 hours prior to harvesting the cells. These stimuli change the transcriptional profile by binding to different receptors with divergent and convergent downstream signaling pathways (**Figure 1B**). All triggers changed expression levels of pre-selected genes that served as positive control for stimulation based on studies with other myeloid cells (such as for example IL-6 for LPS and R848) except TNFα, which was excluded for further analysis (**Figure 1C; Supplemental Figure 1**).

Microglial samples were all subjected to RNA-sequencing using a low input library preparation. After rigorous quality control (**Supplemental Figure 2**), 398 *cultured* microglial samples from 67 different donors and five different brain regions were included (**Supplementary Table 1)**. For details regarding included subjects see **Supplemental Figure 3**. Marker gene expression indicated low levels of contamination of other cells, including neurons, astrocytes, and oligodendrocytes. (**Supplemental Figure 4)**. The percentage of mitochondrial genes detected compared to the total number of expressed genes was only 0.08% and the expression levels of caspases and other apoptotic markers were not found to show a difference in expression across the conditions (**Supplemental Figure 5A**). The receptors for the inflammatory triggers applied in this study are expressed both in *uncultured* and *cultured* conditions, although with some differences (**Supplemental Figure 5B)**. We used expression data from the *cultured* microglial samples as a comparison with stimulated data from the same individuals.

We explored the relative importance of a wide range of biological and technical factors in determining microglial gene expression. As previously observed in *uncultured* microglia (Lopes et al. 2022; Young et al. 2019), we found that technical factors are highly correlated such as proportions of sequenced bases covering mRNA or ribosomal genes (**Supplemental Figure 6A)**, and explained the most variance in expression per gene (**Supplemental Figure 6B)**. We, therefore, controlled for these technical confounders. Using principal component analysis (**Supplemental Figure 7),** we observed no clear separation by stimulation, brain region, sex, and age after regressing out technical factors **(Supplemental Figure 8)**.

### Effects of stimulation on the human microglia transcriptome

To maximize statistical power, we combined samples across multiple brain regions in a linear mixed model accounting for samples of the same donor. We performed differential expression between baseline and each stimulation group separately, controlling for technical factors, brain region, sex, and age. We found widespread differentially expressed genes (DEGs), with the largest number identified in LPS and R848, likely related to power (**Supplemental Figure 9**). Genes with a | log_2_ fold change | > 1 were marked as strongly up or downregulated.

735 genes were significantly upregulated, and 5,723 genes were downregulated after exposure to the TLR4 agonist LPS (**Supplementary Table 2)**. 127 DEGs were strongly upregulated (N = 80 genes) or downregulated (N = 47 genes) (**Figure 2A**). We found 345 up- and 2324 downregulated genes for R848 (**Supplementary Table 3**), a dual TLR7 and TLR8 agonist: 51 (|log_2_ fold change| > 1), and 28 (|log_2_ fold change| < 1) (**Figure 2B**). For both TLR agonists, the strongly upregulated genes were, as expected, significantly enriched for biological pathways related to cytokine signaling and neuroinflammation (**Figure 2A and B**). The strongly downregulated genes were significantly enriched for biological processes such as hormone and amino acid biosynthesis, and cholesterol pathways (**Figure 2A and B)**.

**Figure 2.**
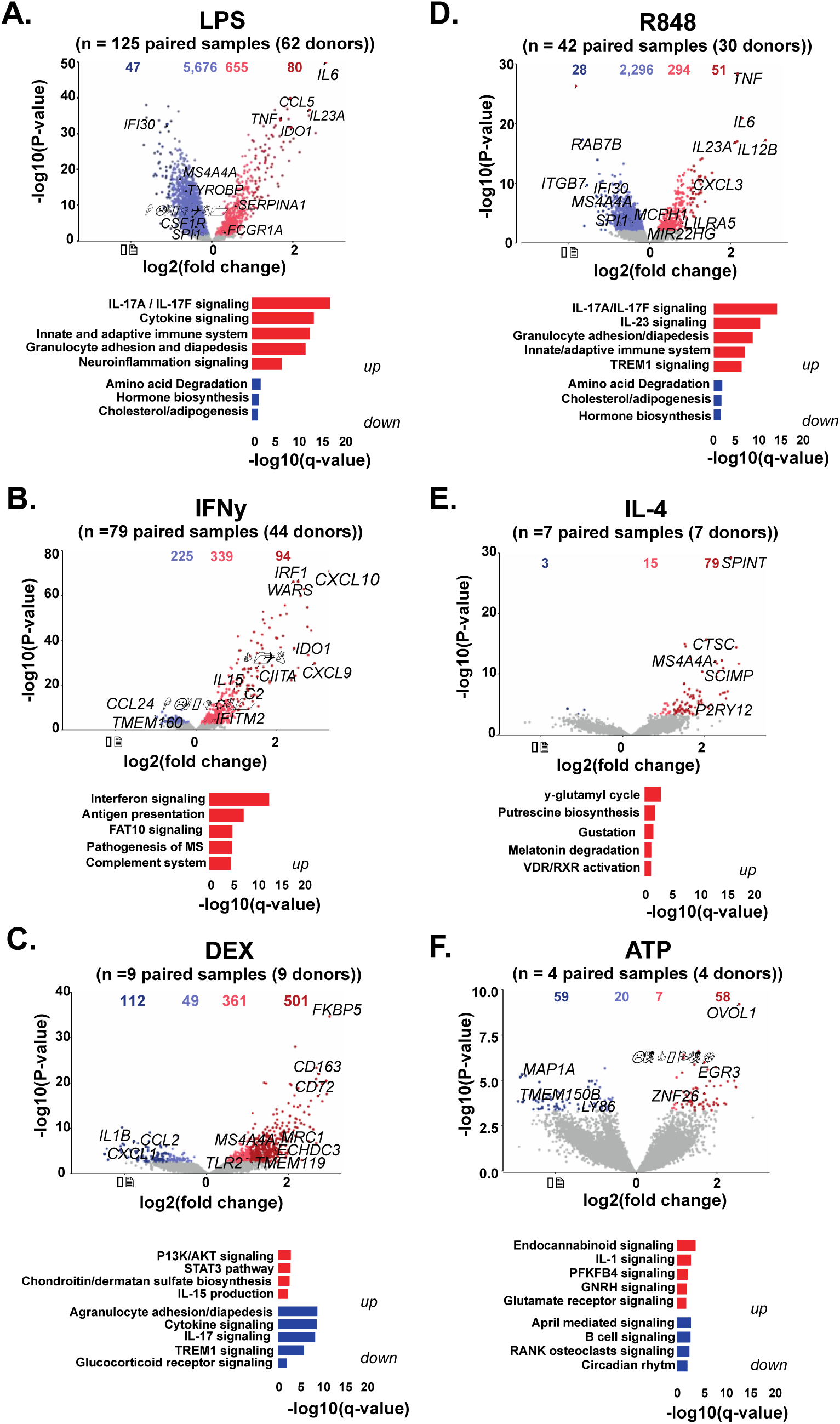
Differential gene expression (DEGs) across stimulations. Volcano plots comparing stimulated microglia to controls in all donors. Genes coloured by whether or not differentially expressed (FDR < 0.05; grey), differentially expressed genes but with modest effects (| log_2_ fold change | 1; orange/blue) and with stronger effects | (log_2_ fold change | > 1; red/dark blue). Numbers of genes in each category above the plot. Well-known microglia specific genes and genes related to Alzheimer’s disease are highlighted. Functional enrichment analyses of all differential expressed genes with log fold change < -1 or > 1 using IPA. Significantly enriched terms are shown with q-value 0.05.

In contrast to LPS and R848, the majority of DEGs were upregulated after IFNγ (**Figure 2C; Supplementary Table 4)** and IL-4 exposure (**Figure 2D; Supplementary Table 5**). The strongly upregulated genes after IFNγ stimulation (N = 94 genes) are mainly components related to IFN signaling and antigen presentation, such as *CXCL10, CXCL9,* and *IRF1 (***Figure 2C**). The strongly upregulated genes after IL-4 stimulation (N = 79 genes) consist of genes related to more general biological processes such as amino acid release and uptake (**Figure 2D).**

Dexamethasone (DEX), a synthetic glucocorticoid, is a potent immunosuppressant. The transcription factor *FKBP5* was 30-fold upregulated. *FKBP5* is an important regulator of stress responses in the brain and acts as a co-chaperone that modulates glucocorticoid receptor activity (Häusl et al. 2021). We found 862 upregulated and 161 downregulated genes after DEX stimulation (**Figure 2E; Supplementary Table 6**). The strongly upregulated genes (N = 501) were enriched for IL-17 and cytokine signaling pathways (**Figure 2E**). The strongly downregulated genes (N = 112 genes) were significantly enriched for gene ontology terms related to neuroinflammation and cytokine signaling and included important genes such as *IL1B* and *CCL2*.

ATP is involved in purinergic signaling and activates microglia causing morphological changes (Inoue, Koizumi, and Ueno 1996). We detected both strongly up (N = 58) and downregulated (N = 59) genes **(Figure 2F; Supplementary Table 7**). Most of these genes were related to general metabolic processes, immune regulation, and controlling biological rhythm **(Figure 2F)**.

Additionally, we used a differential transcript usage (DTU) framework to identify alternative splicing in microglial cells across different stimulations. We identified 779 transcripts in 499 genes with most transcripts coming from the ATP stimulation, suggesting that splicing events increase by ATP-dependent catalyzation. The genes associated with stimulation DTU include genes involved in general biological functions (ribosome, proteosome, antigen presentation and protein binding), but also disease mechanisms, such as atherosclerosis and legionellosis (**Supplemental Figure 10**).

### Unique transcriptional responses across stimulations in human microglia

Next, we compared the transcriptional responses after stimulation between the selected substrates. Correlation of (log_2_) fold change effect sizes between stimulations found strong concordance in direction between LPS and R848 (**Figure 3A**). The DEGs were highly similar between LPS and R848, with the addition of a much higher number of unique DEGs in the LPS stimulation compared to all the other stimulations (**Figure 3B**). We used a threshold of | log_2_ fold change | > 1, to define a ‘*stimulation specific microglia signature*’ comprising 1,006 genes in total that were differentially expressed in one or more conditions (**Figure 3C**). Hierarchical clustering of the stimulations found that LPS, R848 and IFNγ clustered together, separately from IL-4, DEX, and ATP. We observed that the same environmental triggers clustered together in the differential transcript usage framework as for gene expression (**Figure 3D**).

**Figure 3.**
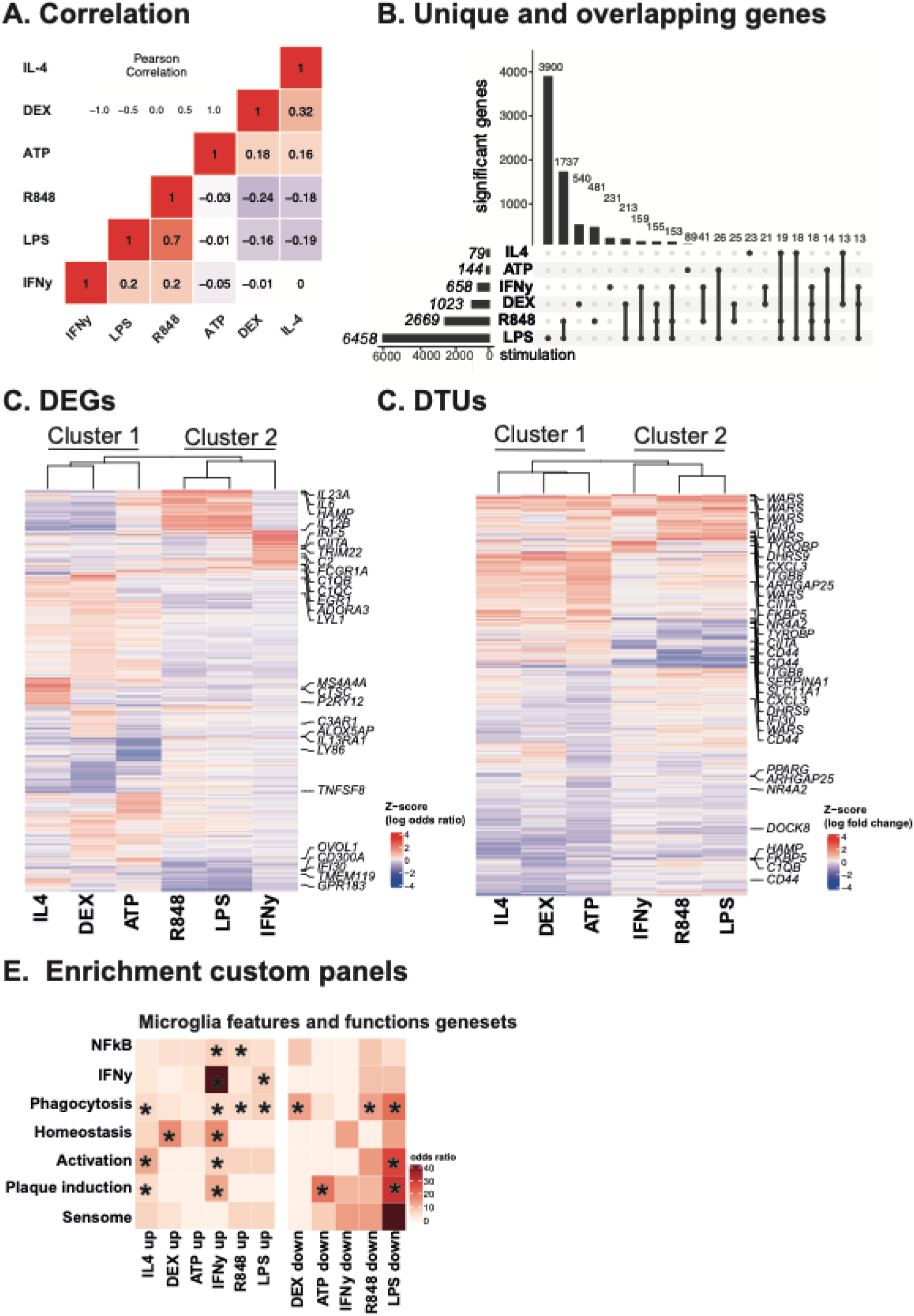
Gene and transcript level expression. A) Pearson correlation values are shown for the pairwise correlations of each stimulation using the log_2_ fold changes for each gene. B) Upset plot depicting number of genes at FDR < 0.05 across all stimulations. C) Heatmap of 1,006 genes significantly changed after stimulation with a Log_2_FC of < -1 or > 1 (FDR 5%). Each gene (row) was plotted as a Z-score of log_2_ fold changes. Well-known microglia and AD-related genes are highlighted. D) Heatmap of 779 transcripts that significantly changed after stimulation with an odds ratio of < -1 or > 1 (FDR 5%). Each transcript (row) was plotted as a Z-score of the median log odds ratio. Well-known microglia and AD related genes are highlighted. E) Enrichment analysis of the top 100 DEGs per stimulation with gene ontology terms related to microglial functions (Grubman et al. 2021; Hickman et al. 2013; Ashburner et al. 2000). Color scheme represents the odds ratio. * Significant P-value from one-sided Fisher exact test.

We performed enrichment analyses of the different stimulation signatures with curated gene ontology terms related to specific microglia functions (Grubman et al. 2021; S. E. Hickman et al. 2013; Ashburner et al. 2000) (**Supplementary Table 8**). Stimulation signatures included well-known microglial markers related to specific microglial functions. The analyses highlighted the involvement of IFNγ signaling in a wide range of other microglial functions besides IFNγ signaling, such as NF-kβ signaling, phagocytosis, homeostasis, and plaque-induced genes (PIGs). The genes that are downregulated after LPS exposure were related to various microglia functions, including phagocytosis, homeostasis, plaque induction and to a unique group of protein transcripts for sensing ligands and microbes, the “microglial sensome” (S. Hickman et al. 2018). Interestingly, the gene ontology term “microglia activation” was enriched only within genes that were downregulated after induction with the TLR-4 agonist LPS. Overall, these results point to the involvement of distinct stimulation signatures, which may play roles in turning on and off specific functions of microglial cells when the environmental context is changed (**Figure 3E**).

### Impact of anatomical brain region on transcriptional responses of microglia after inflammatory stimulation

Although the results presented above assessed the effect of stimulating microglia while controlling for the brain region of origin, we have previously observed differences in gene expression between microglia residing in different regions of the brain, particularly between microglia in the grey matter regions of the neocortex (MFG; STG) with the microglia within the thalamus (THA) and subventricular zone (SVZ) (Lopes et al. 2022). Next, we investigated differences in stimulation response between different brain regions (**Supplemental Figure 11**). We focused only on LPS and IFNγ, the two largest sample sizes. To elucidate whether the pan-brain stimulation results are being driven by a particular brain region, we correlated the log_2_ fold change values of the stimulation results of all brain regions combined (DREAM) with the log_2_ fold change values of each brain region separately. All brain regions positively correlated with the DREAM results, with the strongest correlation seen with SVZ (**Supplemental Figure 12 and 13**). We performed differential expression analyses for each brain region separately and found the largest number of DEGs in the SVZ after LPS and IFNγ compared to baseline (**Figure 4A and B)**. The direction of effect of the DEGs was highly concordant between the brain regions. However, the SVZ microglia showed a consistently stronger response to both LPS and IFNγ stimulation (**Figure 4C and D**), compared to the other brain regions. To further understand the difference in amplitude between SVZ and MFG, we performed gene ontology enrichment analyses for the DEGs in the MFG versus SVZ after LPS and IFNγ stimulation. The SVZ-specific genes (compared to the MFG-specific genes) included genes that were involved in functions related to lipid storage, viral life cycle, hemostasis, and defense response (**Figure 4E and F**).

**Figure 4.**
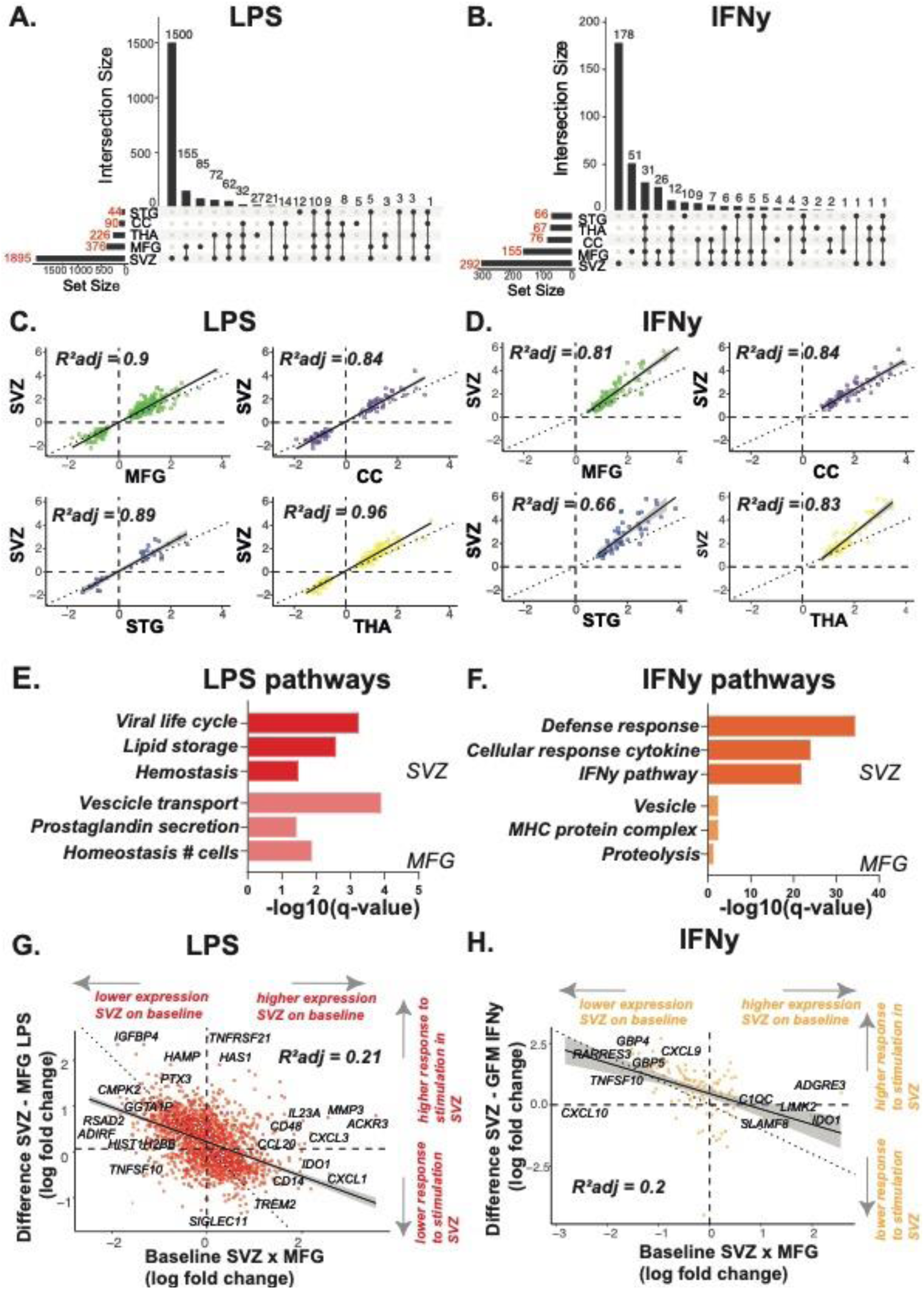
Regional differences after LPS and IFNγ stimulation. A/B) Number of genes differentially expressed with DESeq2 at an FDR < 0.05 in all brain regions separately. C/D) GO biological pathways related to genes that are related to region-specific genes (MFG or SVZ) with | log2 FC | 1. E/F/G) Scatterplot showing the size correlation of LPS or IFNγ response-related genes by brain region. Only genes that were significant in at least one brain region are shown. The x value and y values show the logFC changes of that particular brain region after LPS stimulation. G/H) Scatterplot comparing the difference of the response to stimulation in the SVZ minus the GFM and the expression level of the SVZ times GFM genes on baseline (without stimulation). All differential expressed genes in one of the conditions are shown. MFG: medial frontal gyrus, STG: superior temporal gyrus, THA: thalamus, CC: corpus callosum, SVZ: subventricular zone.

Next, we investigated whether differences in response strength between the MFG and SVZ are related to baseline gene expression. We compared the difference in log_2_ fold change values between these two regions (SVZ minus MFG) after stimulation to the baseline expression of these genes in the unstimulated samples. We found that SVZ genes that are lowly expressed on baseline level compared to the MFG react with a higher amplitude to stimulation. Conversely, genes that are highly expressed on baseline level in the SVZ react less to the stimulation with either LPS or IFNγ (**Figure 4G and H; Supplementary Table 9**). Therefore, region-specific differences in stimulation response strongly depend on basal levels of gene expression.

### Stimulated human microglial cells reveal more overlap with stimulated monocytes than murine microglial cells

We next compared the microglia response to LPS and IFNγ to those from a human monocyte dataset derived from peripheral blood (Navarro et al, in preparation) and response to LPS to those from mouse microglial cells (Gerrits et al. 2020) (**Figure 5A**). Despite having fewer samples per condition, the monocyte dataset had substantially more DEGs per stimulation. We compared the log_2_ fold change of the monocyte DEGs with those in our microglia. We observed 2-3 times higher log_2_ fold change changes for monocytes compared to microglia. Both these results point towards stronger responses of isolated human monocytes compared to isolated human microglia. To counter the differences in effect strength, for monocytes, we used a gene list of DEGs with a threshold of | log_2_ fold change | > 5 after LPS or IFNγ stimulation for 24 hours. We found that the signature gene lists for monocytes strongly overlapped with microglia for the upregulated and downregulated genes after LPS exposure, but only with genes upregulated after IFNγ exposure (**Figure 5B**). We identified in total 4,628 genes that are differentially expressed in both conditions (monocytes and microglia) for LPS and 534 genes for IFNγ and identified around 4-13% genes with opposite directions of effect (adjusted R^2^ = 0.62). A few stimulation-specific LPS genes (e.g. *C5AR1, NR4A2, RAB20, CD163*) are downregulated in microglia but upregulated in monocytes after LPS exposure (adjusted R^2^ = 0.44). *HAMP* expression, for example, is increased in microglia but decreased in monocytes after LPS exposure. Only a few IFNγ specific genes showed opposite direction of effect (e.g. *FES, DUSP23, CREG1, TMEM138)* in microglia compared to monocytes (**Figure 5C and D**).

**Figure 5.**
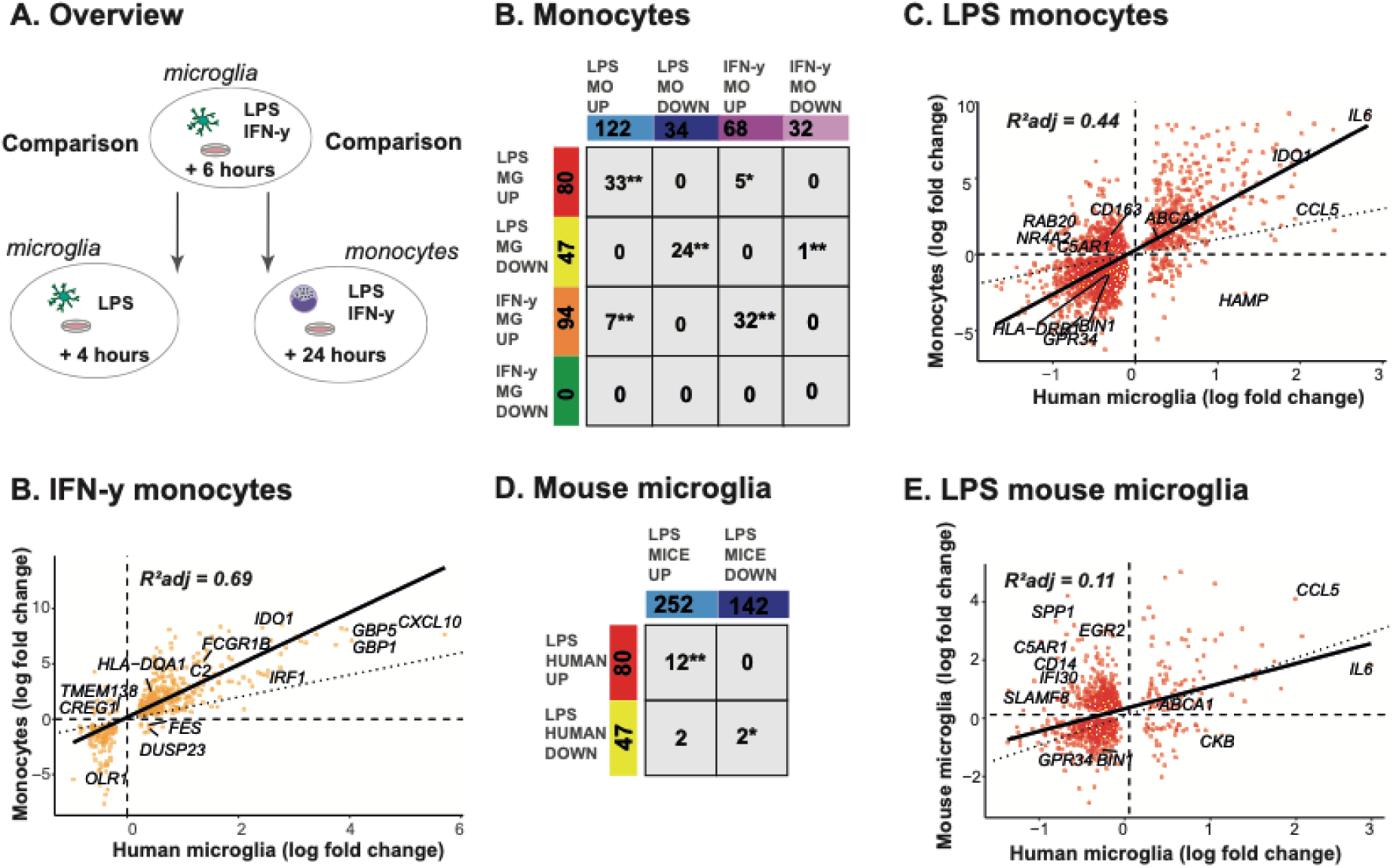
Comparison between LPS stimulated human microglia and human monocyte and mouse microglia data. A) Overview of comparisons. B) Replication analysis with an independent dataset of cultured monocytes and stimulated with LPS and IFNγ for 24 hours, 35 samples in total. Only strongly up- and downregulated genes are compared between cultured monocytes (|log2FC| > 5 and human microglia (|log2FC| > 1). Number of overlapping genes between two conditions are presented. The asterisk indicates significant enrichment by a one-sided Fisher’s exact test. B) Scatterplot showing the size correlation of LPS and IFNγ response genes in human microglia and monocytes. The x-axis shows the logFC values in human microglia and the Y-axis shows the logFC in the cultured monocytes. Only genes that were significant (FDR < 0.05) in both conditions shown. D) Replication analysis with an independent dataset of mice microglial cells stimulated with LPS for 4 hours, 3 samples in total. Only strongly up and downregulated genes are compared between mice microglia (|log2FC| > 1 and human microglia (|log2FC| > 1). Number of overlapping genes between two conditions are presented. The asterisk indicates significant enrichment by a one-sided Fisher’s exact test. E) Scatterplot showing the size correlation of LPS response genes in human microglia mice. The x-axis shows the logFC values in human microglia and the Y-axis shows the logFC in mouse microglial cells. Only genes that were significant (FDR < 0.05) in both conditions are shown.

To explore the differences between functional profiles in human and mouse microglia data, we also compared our LPS signature gene list with LPS stimulated microglial cells in mice (Gerrits et al. 2020). We found a small but significant enrichment between the LPS up-respectively downregulated genes in human and mice microglia (**Figure 5E**). We identified 1,258 genes that are differentially expressed in both conditions (human and mouse microglia) and the correlation was positive (**Figure 5F**). However, around 40% of the shared significant genes were in the opposite direction of effects (adjusted R^2^ = 0.11). Well-known specific stimulation markers such as *IL6, CCL5* were upregulated in both human and murine microglia after LPS exposure. Well-known microglia genes *BIN1* and *GPR34,* for example, were downregulated in both human and mouse microglia after LPS exposure. However, most downregulated genes in human microglia were upregulated in the microglia of mice after LPS stimulation (e.g. *C5AR1, IFI30, ITGB7, SLAMF8, SPP1, CD14, EGR2)*. (**Figure 5D**). In contrast to the monocytes, the amplitudes were in the same range, and for instance, *IL6* responded stronger in human microglia.

### Application of the atlas to show genetic interactions with stimulation in the human microglia transcriptome

Next, we integrated our gene expression profiles with common genetic variants to map *cis*-expression QTLs (eQTLs), a set of statistical associations between a single nucleotide polymorphism (SNP) and the expression of a nearby gene. We performed *cis*-eQTL analyses in primary human microglia that are cultured and stimulated with LPS or IFNγ. We selected two brain regions with the largest number of donors for these analyses (MFG and SVZ). After QC, genotypes from 64 individuals of European ancestry were used for the analysis (**Supplemental Figure 14 and 15**). However, due to uneven sampling, the individual brain region cohorts ranged from 21 to 36 unique donors. Due to the low number of donors in each individual cohort, only 1 cis-eQTL was found at a permissive false discovery rate of 20%.

Next, we investigated the possibility of using the data to map response *cis*-eQTLs (reQTLs), whereby there is a statistical interaction between genotype and the effect of stimulation on the expression of a gene. This has been observed in other myeloid cells but not in microglia. We used the suez framework (Knowles et al. 2018) to detect reQTLs for either LPS or IFNγ stimulation, combining all available samples across brain regions. Suez accounts for both mixed ancestries and overlapping donors using a linear mixed model.

For mapping reQTLs to LPS stimulation, we used 255 samples from 66 unique donors. Given the small sample size, we did not find any reQTLs at a false discovery rate of 5% but identified 9 genes with a reQTL at an FDR of 20%. (**Supplemental Figure 16; Supplementary Table 10**). The top reQTL by P-value was seen in *HLA-DQA1*, where LPS stimulation reduces the strength of the association of the rs3104367 genotype with *HLA-DQA1* expression (**Figure 6A**; q-value = 0.0038). Additionally, an LPS reQTL was found in *DNAAF1* (**Figure 6B**; q-value = 0.0357). Of the 9 LPS reQTLs, only *DNAAF1* was differentially expressed by LPS treatment (LFC = 1.33; adjusted P < 1e-16). The *DNAAF1* reQTL acts to reduce the strength of association between rs78885610 and *DNAAF1* in LPS stimulation compared to baseline.

**Figure 6.**
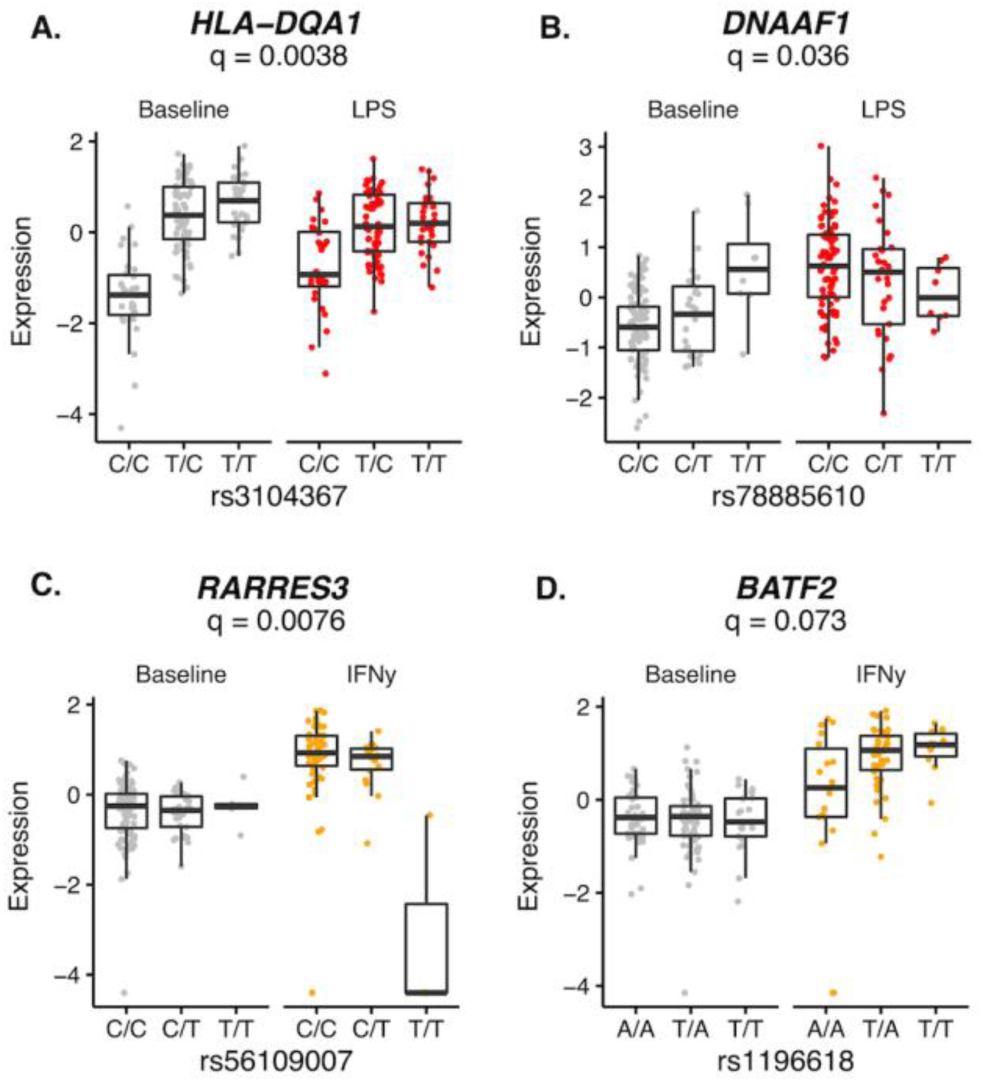
Mapping response cis-eQTLs (reQTLs) in microglia. A-D) Boxplots showing correlation with gene expression variation in stimulated conditions, indicating gene-environment interactions. q refers to adjusted P-value following correction for numbers of tested variants and genes.

For IFNγ stimulation, we used 211 samples from 64 unique donors and identified 6 genes with a reQTL at FDR 20% (**Supplemental Figure 17; Supplementary Table 10)**. Of the 6 reQTL genes, 2 were differentially expressed by IFNγ stimulation: *RARRES3* (**Figure 6C**; q-value = 0.0076; LFC = 2.84; adjusted P < 1e-16) and *BATF2* (**Figure 6D**; q-value = 0.073; LFC = 2.77; adjusted P < 1e-16)

### Defining the association of stimulation signatures in human microglia with aging and Alzheimer’s disease and aging

To illustrate some examples of how this atlas can be applied, we used it to assess the effect of aging on the cultured microglial cells by fitting a linear mixed model accounting for shared donors across all brain regions. We observed 311 genes (262 up, 49 down at FDR < 0.05) associated with chronological age of subjects in these samples (**Supplementary Table 11**). The upregulated genes in chronological aging showed overrepresentation for genes that are upregulated after LPS stimulation, but not after R848 stimulation. Downregulated genes after LPS stimulation were also significantly enriched in genes that were also downregulated in aging. Important LPS response genes that are associated with chronological aging are *AIM2, IL15, VNN2, ZNF23* (**Supplemental Figure 18**). Also, well-known DEX stimulation response genes were significantly enriched for genes associated with chronological aging, including *NR3C1, CD163,* and *PFKP (***Supplemental Figure 18**).

Next, we performed enrichment analyses of the different stimulation signatures with curated gene lists for brain diseases based on TWAS or eQTL datasets (Raj et al. 2018; Lopes et al. 2022; Y. I. Li et al. 2019). The top 100 upregulated genes after DEX stimulation showed enrichment with disease risk genes for Alzheimer’s disease (AD) (*BIN1, PTK2B*), and genes that were upregulated after IFNγ stimulation with Multiple Sclerosis (MS) (*CIITA, IRF5, STAT2)*. In contrast, the top 100 downregulated genes after R848 stimulation showed enrichment with disease risk genes for Parkinson’s disease (PD) (*HSD3B7, NCKIPSD, PTPN22*) (**Figure 7A; Supplementary Table 12**). Next, we focused specifically on genes associated with AD risk through TWAS (Raj et al. 2018) (**Figure 7B**), GWAS (Bellenguez et al. 2022), and microglia-specific genes that previously have been associated with AD pathology in eQTL datasets (Lopes et al. 2022) (**Figure 7C**). For example, the GWAS-annotated AD risk genes *MS4A4A* and *BIN1* are significantly upregulated after DEX and IL-4 stimulation, but significantly downregulated after LPS and R848 exposure (**Figure 7C**). We also analyzed changes in splicing and gene expression levels of several microglia-specific genes in mechanistic AD studies (Lu et al. 2022) across stimulations. By analyzing the differential transcript usage (DTU) framework, we identified changes in splicing, such as *TYROBP* in LPS (**Supplemental Figure 19**), *WARS* (**Figure 7D**) *CIITA* (**Supplemental Figure 19**) in IFNγ, *WDR74* in R848, and *PPARG* in IL-4 stimulated samples compared to the unstimulated microglia samples. We also analyzed the differential gene expression of interesting microglia-specific genes previously associated with immunoregulatory mechanisms in AD pathology. For example, we show two microglia-specific genes associated with AD pathology in mechanistic studies (Nitsch et al. 2021; Guillemin et al. 2005). *IL23A* was significantly upregulated after LPS or R848 stimulation and downregulated after exposure to DEX and IL-4. Indoleamine 2,3 dioxygenase 1 (*IDO1)* was significantly upregulated after stimulation with LPS, IFNγ or R848 and downregulated after stimulation with IL-4 (**Figure 7E**).

**Figure 7.**
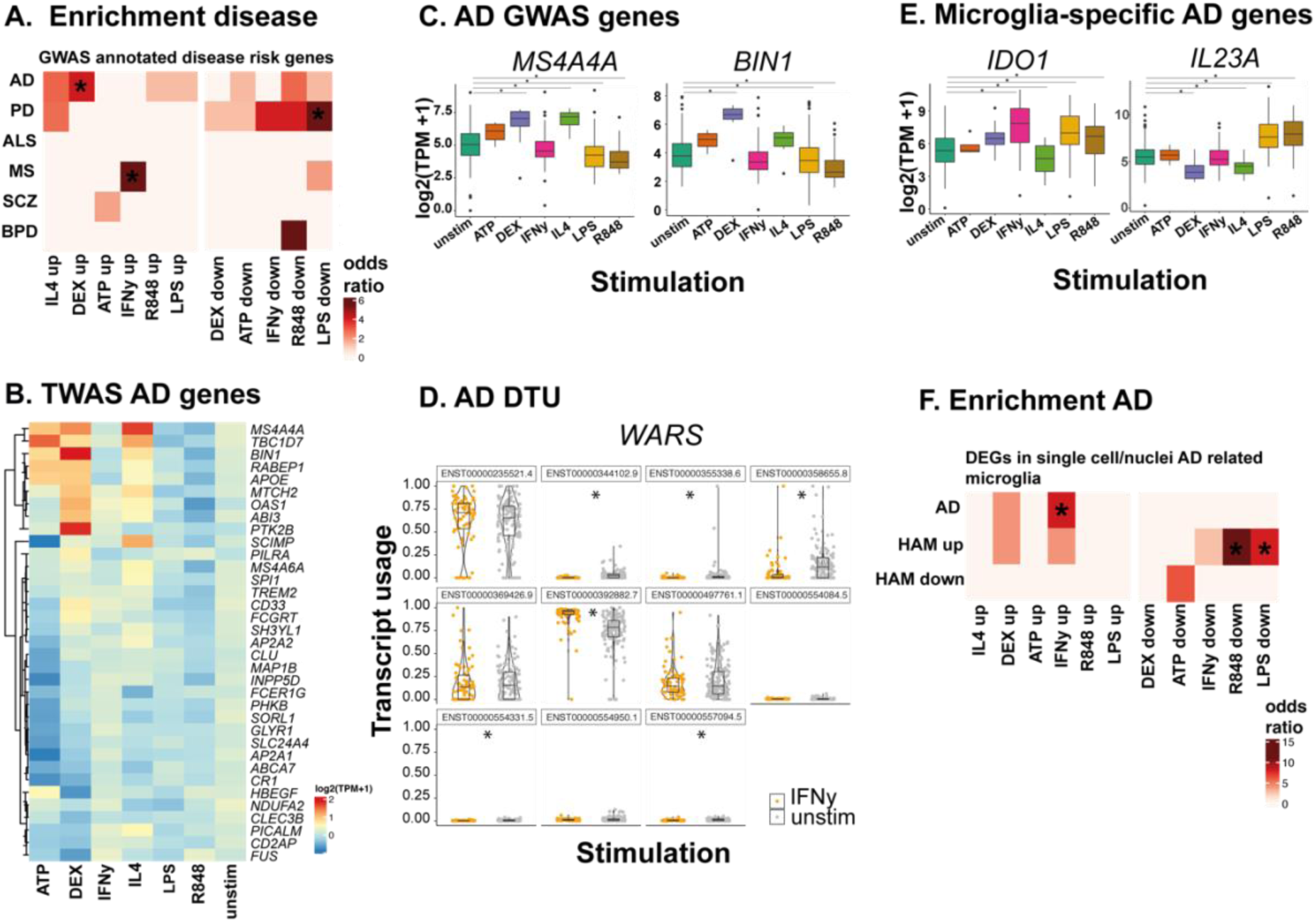
Defining the association between stimulation signature genes and Alzheimer’s disease. A) Enrichment analysis of the top 100 DEGsper stimulation with disease related genes based on TWAS and eQTLs, Color scheme represents the odds ratio. * Significant P-value from one-sided Fisher exact test. B) Expression levels (TPM+1 log 2 scale) of well-known Alzheimer disease (AD) genes across different stimulations (unstimulated (unstim), Lipopolysaccharide (LPS), Interferon-gamma (IFNγ), Tumor necrosis factor alpha (TNFα), Resiquimod (R848), Adenotriphosphaat (ATP), Dexamethasone (DEX), Interleukine-4 (IL-4), for 6 hours. Blue colors indicate low expression and red colors indicate high gene expression.C) Examples of GWAS annotated AD disease risk genes DEGs in each stimulation. P-values two-sided are FDR adjusted from the linear mixed model of the differential expression analysis. The box plots show the median, the box spans the first to third quartile and the whiskers extend 1.5 times the IQR from the box. D) Example transcript usage of WARS in IFNγ. The DTU signal is driven by an increase resp. decrease in the transcripts ENST0000392882.7, ENST00000344102.9, ENST00000554331.5, ENST00000557094.5, ENST0000358655.8, ENST00000355338.6. E) Examples of microglia-specific genes associated with AD pathology DEGs in each stimulation. P-values two-sided are FDR adjusted from the linear mixed model of the differential expression analysis. The box plots show the median, the box spans the first to third quartile and the whiskers extend 1.5 times the IQR from the box. F) Enrichment analysis of the top 100 DEGs per stimulation with disease related genes based on TWAS and eQTLs, gene ontology terms related to microglial functions, and single cell and nuclei data related to AD pathology. Color scheme represents the odds ratio. * Significant P-value from one-sided Fisher exact test.

Next, we compared the stimulation signature gene set with specific single cell/nuclei microglia gene lists related to AD pathology (**Figure 7F; Supplementary Table 13**). We found that single nuclei microglia of AD brains (Mathys et al. 2019) are significantly enriched for genes that are upregulated after IFNγ stimulation. The genes downregulated after LPS and R848 exposure showed enrichment with genes that are upregulated in the postmortem human Alzheimer’s brain (Srinivasan et al. 2020).

## Discussion

In the present study, we describe MiGASti, a comprehensive resource of transcriptomic and genetic data of human microglial cells cultured and exposed to different stimuli. For this study, isolated microglial cells were exposed to a range of triggers to create the largest study so far on human microglial transcriptional responses to environmental challenges. This has allowed us to identify divergent patterns of differentially expressed genes and transcripts in response to stimulation, and to identify stimulation specific genes associated with biological processes, functions, and diseases, such as aging and Alzheimer’s disease. This atlas enabled us to define distinct inflammatory responses in microglia of different anatomical brain regions. This study also provides insight into differences between functional profiles of human microglia in comparison to human monocytes and mouse microglia. By integrating common genetic variants, we were able to map response cis-eQTLS (reQTLs), showing interactions between genotypes and the effect of LPS or IFNγ stimulation on the expression of a gene. We believe this resource will support further investigations into the contribution of human microglia states to health and disease.

Clustering the stimulation signature gene lists shows similarities and differences between the transcriptional response to the six triggers of interest of this study. In line with previous studies in primary human microglia (J. Melief et al. 2016), human macrophages (Cheng et al. 2019; Alasoo et al. 2015; Xue et al. 2014; Emam et al. 2020), and murine microglia (Holtman et al. 2015; Lund et al. 2006; Das et al. 2018; Lively and Schlichter 2018), we found clustering of the responsome to triggers often referred to as (pro-) inflammatory, including LPS, R848 or IFN, and the triggers that are classically viewed as ‘restorative’, such as IL-4 and dexamethasone. The response to LPS and R848 showed a large overlap, which could be expected as the MyD88-dependent signaling cascades following binding of these factors to their Toll-like receptors converges on the activation of NF-kB (Kawai and Akira 2007). ATP, which is often seen also as an (pro-) inflammatory trigger, clustered with the response to IL-4 and dexamethasone. Of note, we used high concentrations of ATP (1 mM), known to stimulate the P2X7 receptor (Calovi, Mut-Arbona, and Sperlágh 2019). Triggering of P2X7 receptors with ATP, has been shown to lead to release of neurotrophic factors and the activation of the NOD-, LRR- and pyrin domain-containing protein 3 (NLRP3) inflammasome (Campagno and Mitchell 2021). Although purinergic signaling has also been linked to NF-kβ activation (Suzuki et al. 2020), our results suggest that this was not the case in our stimulation regimen, as we did not see clustering with the other NF-kβ stimulating compounds. Our study where we show different transcriptional responses to a range of triggers is in line with a recent consensus paper (Paolicelli et al. 2022). Microglia cannot be seen in dichotomic phases, separating them between active or inactive, as M1 or M2-stimulated, or as stimulated by pro- or anti-inflammatory triggers. The profiles that we generated contribute to the view that microglia exist in diverse, dynamic, and multidimensional states depending on the context, including their local environment (Paolicelli et al. 2022). Even in an artificial in vitro situation, where the local environment can be skewed in a controlled matter, this heterogeneity was clearly observed.

We compared the transcriptomic responsome of human microglia to the response of human monocytes, a more accessible cell-type for research, but also a cell-type that has the potential as an application for biomarker discovery. Interestingly, we observed that the amplitudes of the response in monocytes were generally higher compared to human and murine microglia. These results are in line with a previous study, where we compared LPS responses between primary microglia, monocytes, monocyte-derived macrophages, and monocyte-derived dendritic cells in the same study (J. Melief et al. 2016). Monocytes and monocytes differentiated to dendritic cells and macrophages responded much stronger to LPS than microglia. Microglia are often viewed as an immune cell type which is less responsive compared to other myeloid cell subsets to protect the brain from unnecessary injury due to inflammatory processes (Spiteri et al. 2023). Monocytes on the other hand, are involved in first line defense, and are generally considered more efficient in quick clearance, in comparison to microglia cells (Fani Maleki and Rivest 2019). Our results are in line with this view. However, technical factors provide an alternative explanation for these results. Monocytes can be readily isolated from blood, whereas for microglia longer and harsher methods are required, resulting in some level of reactivation of the cells. A study in mice where *in vivo* LPS responses were compared to *ex vivo* responses suggest that isolation does not dampen the response (Lund et al. 2006). Another study in rodents show that differential responses between myeloid cell subtypes might be trigger/pathway dependent (Janova et al. 2016). Our results comparing SVZ and MFG show that baseline expression levels dictate the amplitude of the transcriptional response to in vitro stimulation. By comparing our human microglia LPS signatures to mice, we even found most gene expression changes go in the opposite direction, suggesting that LPS responses of murine microglia are not representative for human microglia.

Mechanistic studies on the role of microglia in pathology often focus on genes and pathways which are upregulated after inflammatory triggers, such as LPS and IFN. Our results suggest it might be equally important to study genes and pathways that are downregulated when the inflammatory context of the cells is changed, or genes and pathways which are upregulated in the homeostatic/restorative context. We did not detect a significant association between AD-related genes and genes upregulated after LPS, R848 or IFNγ. Instead, we found enrichment of these genes in cells stimulated with DEX or IL-4, as well as in genes that are downregulated after LPS, R848 or IFN©. This view is in line with previous studies in primary murine microglia that showed genes downregulated by LPS/IFN© treatment microglia were involved in neurodegenerative disease (Pulido-Salgado et al. 2018). Downregulated genes due to pro-inflammatory treatments are often unnoticed and involvement in inflammation has been questioned. Importantly, our results of up- and downregulated genes after skewing human microglia phenotypes *in vitro* are relative results where we compare unstimulated with stimulated cells. The results can also be explained as follows: AD-associated genes are enriched in upregulated genes of unstimulated versus LPS/IFNγ- stimulated states. Disease risk may therefore play a role in functions which are repressed when microglia are skewed by the more classical inflammatory triggers, or functions upregulated in homeostatic/restorative conditions. Interestingly, we found that downregulated genes after LPS and R848 stimulation in human microglia have been associated with the genes upregulated in human Alzheimer’s brains. We did not find enrichment of upregulated genes after LPS or R848 stimulation in downregulated genes in Alzheimer’s brains. These results therefore suggest a specific dysregulation of pathways that are downregulated in response to inflammatory triggers. Genes downregulated by these inflammatory stimuli have many important microglial functions, such as phagocytosis, plaque induction, sensome ligand scanning (**Figure 3B**) and are involved in many microglial functions that are disrupted in AD brains (with signs of microglia activation) (**Figure 7E**). Downregulation of these genes may contribute to AD risk, consistent with earlier findings in AD (McQuade and Blurton-Jones 2019; Solomon et al. 2022; C.-C. Liu et al. 2013). More in detail, LPS and R848 decrease expression of many AD related genes in the same direction as how disease related genetic variants change gene expression (see **Table 1).** This suggests that LPS/R848-like triggers can induce a similar phenotype as what we know contributes to the disease through genetic variation related to AD and a second hit like a viral infection might contribute to a genetically AD-prone microglia phenotype.

**Table 1.** AD genetic risk variants and the direction of stimulation.

| Gene | Significant effects of stimulation (direction) | Proposed microglial-related function in AD |
| --- | --- | --- |
| *MS4A4A* | **LPS (↓), R848 (↓)**, IL-4 (↑), DEX (↑) | Transmembrane protein that interacts with TREM2, thought to modulate inflammatory responses and function as a lipid receptor. The protective variant increases MS4A4A expression and shifts the 'chemokine microglia subpopulation' to an interferon state, while the risk variant suppresses MS4A4A expression. The protective variant impacts a specific population of microglia in human brains that regulates cytokines, chemokines, and the Type-I interferon pathway (You et al. 2023) |
| *OAS1* | IFNγ (↑), **R848 (↓)** | The risk allele is associated with decreased OAS1 expression. OAS1 is part of an IFN-γ signaling network and significantly upregulated in AD brain tissue. In vitro modelling shows that decreased OAS1 is associated with exaggerated production of TNF-α with IFN-γ stimulation (Magusali et al. 2021) |
| *SCIMP* | IL-4 (↑), DEX (↑) | Alteration in TLR mediated microglial phagocytosis via MHC II (Vogrinc, Goričar, and Dolžan 2021) |
| *BIN1* | **LPS (↓), R848 (↓)**, DEX (↑) | Risk variants decrease BIN1 enhancer activity and, in turn, BIN1 expression, specifically in microglia and other myeloid cells. Decreased expression of BIN1 in microglia impairs phagocytosis and inflammatory response, including the response to LPS (Sudwarts et al. 2022) |
| *TREM2* | **LPS (↓), R848 (↓)** | Cell-surface receptor binding a variety of ligands including APOE. Triggering of TREM2 leads to chemotaxis, proliferation, survival and phagocytosis in microglia. Risk variants impair the function of TREM2 (Podlešny-Drabiniok, Marcora, and Goate 2020) |
| *APOE* | **R848 (↓)** | Present in amyloid -β and contributes to clearance and amyloid deposition. APOE variants related to increased AD risk reduce amyloid -β clearance (Shi and Holtzman 2018) |
| *CD33* | LPS (↓), R848 (↓) | Disease variants are related to splicing of CD33. Disease-related CD33 isoforms repress phagocytosis in microglia, whereas the protective variants shows enhanced phagocytosis and uptake of amyloid -β (Eskandari-Sedighi, Jung, and Macauley 2023) |

Red/bold indicates effects of stimulation on gene expression in line with the effects of genetic variants that increase risk to AD.

Following our work on the Microglia Genomic Atlas (MiGA) (Lopes et al. 2022) where we mapped expression QTLs in *ex vivo* microglia, we made the first effort to map response eQTLs in microglia to LPS and interferon. We aimed to provide a useful resource for evaluating the genetic explanation for inter-individual variation to immune responses.

### Limitations

One of the limitations of these *in vitro* experiments with *cultured* microglial cells is that they cannot capture situations in an *in vivo* setting. First, we isolated microglia from the brain. Besides neuroinflammation, hypoxia, and long postmortem intervals, technical artifacts (enzymatic digestion, temperature changes, sorting) may cause microglial activation. Microglia isolation from their environment as well as culturing them alters their function and will lead eventually to a partial loss of their signature gene expression (Gosselin et al. 2017). We, therefore, put the cells in culture for a limited amount of time before we stimulated the cells. Second, our stimulations do not reflect the *in vivo* situation during an insult or change in the cellular environment. This will lead to a cascade of production of many other soluble and cell-bound signaling molecules, by microglia and other cell types of the brain, which regulate the full response to this trigger in time. These factors will also alter gene expression in microglia. We investigated and compared transcriptional responses of different stimuli after 6 hours of stimulation, because this time point offers a good picture of the first wave of transcriptional changes, before other cellular factors start to play a major impact. Also, we did not detect a clear separation using PCA by stimulation or other factors likely due to the high donor-to-donor variation in microglial cells. We were dependent on the available data for comparisons with monocyte and microglia mouse data, even though stimulation incubation times were not equal. Overall, the sample sizes are the largest thus far, but still very small for some of the stimulations, as for example, ATP. Also, the low number of response eQTLs detected in the interaction analyses is likely due to the small sample size per stimulation and/or high donor-to-donor variation in microglial cells. Our low discovery rate, even at a permissive FDR, is likely due to both low sample sizes but also to the relatively weak response to stimulation, compared to monocytes, and a higher inter-donor variation. While this may be improved with increasing sample size, we also look to single-cell efforts to define clusters of microglia in particular states, to map QTLs there.

Taken together, our data provide a comprehensive database of gene expression alterations and genetic effects in microglial cells after exposure to different stimuli. Our data are a useful resource for the interpretation of current data but also for the design of future experiments.

## Methods

### Human brain tissue

Post-mortem brain samples were obtained from the Netherlands Brain Bank (NBB). The permission to collect human brain material was obtained from the Ethical Committee of the VU University Medical Center, Amsterdam, The Netherlands. Informed consent for autopsy, the use of brain tissue and accompanied clinical information for research purposes was obtained per donor ante-mortem. Neuropathological assessments have been performed by the NBB (see URLs). Detailed information per donor, including tissue type, age, sex, postmortem interval, pH of cerebrospinal fluid, cause of death, diagnosis, use of medication and neuropathological information is provided in **Supplementary Table 14**.

### Microglial isolation

Brain tissue was stored in Hibernate media (Gibco) at 4 °C upon processing within 24 hours after autopsy. Microglia were isolated from six regions, including medial frontal gyrus (MFG; 43), superior frontal gyrus (SFG; 1 samples), superior temporal gyrus (STG; 34 samples), thalamus (THA; 24 samples), subventricular zone (SVZ; 32 samples) and corpus callosum (CC; 16 samples). Microglia were isolated as described before in detail(J. Melief et al. 2016; Böttcher et al. 2019; Sneeboer et al. 2019). In brief, brain tissue was first mechanically dissociated through a metal sieve in a glucose- potassium-sodium buffer (GKN-BSA; 8.0 g/L NaCl, 0.4 g/L KCl, 1.77 g/L Na2HPO4.2H2O, 0.69 g/L NaH2PO4.H2O, 2.0 g/L D-(1)-glucose, 0.3% bovine serum albumin (BSA, Merck, Darmstadt, Germany); pH 7.4) and supplemented with collagenase Type I (3700 units/mL; Worthington, USA) and DNase I (200 µg/mL; Roche, Switzerland) or 2% of Trypsin (Invitrogen) at 37 °C for 30 min or 60 min while shaking. The suspension was put over a 100 µM cell strainer and washed with GKN-BSA buffer in the centrifuge (1800 rpm, slow brake, 4 °C, 10 min) before the pellet was resuspended in 20 mL GKN-BSA buffer. 10 mL of Percoll (Merck, Darmstadt, Germany) was added dropwise and the tissue homogenate was centrifuged at 4000 rpm (fast acceleration, slow brake at 4 °C, 30 min). The middle layer was collected and washed with GKN-BSA buffer, followed by resuspension, and centrifuging in a magnetic-activated cell sorting (MACS) buffer (PBS, 1% heat-inactivated fetal cow serum (FCS), 2 mM EDTA; 1500 rpm, 10 °C, 10 min). Microglia were positively selected with CD11b-conjugated magnetic microbeads (Miltenyi Biotec, Germany) according to the manufacturer’s protocol. This protocol has been validated and the resulting cell viability was between 70 and 98% (Jeroen Melief et al. 2012). We have previously shown by mass cytometry and single-cell RNA seq that the levels of infiltration of macrophages and monocytes, and other brain cells is low (Lopes et al. 2022; J. Melief et al. 2016; Sneeboer et al. 2019).

### Culturing and stimulation of microglial cells

Microglia were cultured in a poly-L-lysine (PLL; Merck, Germany) coated 96-wells flat bottom plate (Greiner Bio-One, Austria) at a density of 1.0 × 10^5^ cells in a total volume of 200 μL Rosswell-Park-Memorial-Institute medium (RPMI; Gibco Life Technologies, USA) supplemented with 10% FCS, 2 mM L-glutamine (Gibco Life Technologies, USA), 1% penicillin–streptomycin (Gibco Life Technologies, USA) and 100 ng/ml IL-34 (Miltenyi Biotech, Germany). After overnight incubation, pMG were stimulated with 100 ng/mL lipopolysaccharide (LPS) from Escherichia coli 0111:B4 (Merck, Germany), or with 50 ng/ml interferon gamma INF-γ (Peprotech, London, UK), 1ug/ml Resiquimod (R848), 50 ng/ml tumor necrosis factor alpha (Sigma-Aldrich, The Netherlands), 40 ng/ml IL-4 (Peprotech, London, UK), 1000 nM dexamethasone (Sigma-Aldrich, The Netherlands), or 1 mM ATP (Sigma-Aldrich, The Netherlands) for 6 h. The concentrations and incubation times for LPS and IFNg are based on dose-response curves and 6 hours showed the highest fold change of expression of marker genes. We added the other stimulations as comparison. The cells were harvested with TRIzol reagent and stored at -80 Celsius degrees for further analysis. Microglial RNA was isolated using the TRIzol method (Rio et al. 2010) and cDNA libraries were generated Genewiz using the Ultra-low input system which uses Poly-A selection. SMART-Seq v4 Ultra Low Input RNA Kit was used for library construction using < 100 ng of RNA. The libraries were sequenced as 150 bp on fragments with an average read depth of 28 million (ranging from 0.06-128M) read pairs on the Illumina HiSeq 2500. RNA-seq data was processed using the RAPiD pipeline (Wang et al. 2015). RAPiD aligns samples to the hg38 genome build using STAR (Dobin et al. 2013) using the GENCODE v30 transcriptome reference and calculates quality control metrics using Picard (Toolkit 2016). RNA-seq quality control was performed applying four filters to remove samples: 1) samples with less than 1 M reads aligned from STAR; 2) samples with more than 20% of the reads aligned to ribosomal regions; 3) samples with less than 5% of the reads mapping to mRNA; 4) samples with high chance of RNA degradation based on gene coverage plot. Estimated transcript abundance was obtained using RSEM (B. Li and Dewey 2011) and transcripts were summed to the gene level with tximport (Love, Soneson, and Robinson 2017). Genes with more than 1 read count per million (CPM) in 50% of the samples were kept for downstream analysis. Gene level read counts were normalized as transcripts per million mapped reads (TPM) to adjust for sequencing library size differences. We used several approaches to test the quality of the included samples: 1) the percentage of mitochondrial genes was only 0.87% on average 2) the expression of several apoptotic markers, such as *CASP3, BTG1* was low across all stimulations 3) the expression levels of several well-known specific stimulation markers showed up- and downregulation patterns as expected (Figure 1C and Supplemental Figure 1).

### DNA isolation and genotyping

Genomic DNA was extracted from brain sections using the Qiagen DNeasy Blood & Tissue Kit and followed the manufacturer’s instructions. Briefly, a small piece of brain tissue was cut and placed in a 96-deepwell plate (Thermo Fisher) on dry ice. A mastermix containing binding enhancer, PBS and proteinase K (Qiagen) was added to each well. The solution was incubated overnight at 65 degrees Celsius. The KingFisher™ (KF) Duo system was applied with a 12 pin magnet head which enabled processing of 12 samples per run using microtiter 96 deepwell plates (Thermo Fisher). Prior to the extraction process, the deepwell plates were filled with the following reagents: wash buffer (Qiagen), 80% Ethanol, elution buffer (Qiagen) and tip combs (Qiagen). For extraction, 440 μl of DNA binding buffer beads (Qiagen) were added to the sample and mixed by pipetting. After that, automated extraction was executed and completed within 30 minutes using the protocol provided by the manufacturer. The extracted DNA was eluted in 200 μl elution buffer and subsequently transferred to 1.5 ml tubes for storage. DNA quality and concentration was assessed using a Nanodrop. Samples were genotyped using the Illumina Infinium Global Screening Array (GSA), which contains a genome-wide backbone of 642,824 common variants plus custom disease SNP content (∼ 60,000 SNPs).

### Sources of transcriptomic variation

To understand major sources of variation in the gene expression data at the sample level, we used PCA and linear regression to measure the effect of the following experimental confounders on gene expression variance: sex, age, donor, brain region, stimulation, and technical covariates estimated by Picard. We then applied variancePartition (v1.17.7), which uses a linear mixed model to attribute a percentage of variation in expression based on selected covariates on each gene. Gene counts were normalized using trimmed means of M-values. (TMM) values calculated from edgeR and voom transformed, which is a method that estimates the mean-variance relationship of the log-counts as input to variancePartition. The technical covariates included in the analysis were % mRNA bases (Picard), % ribosomal bases (Picard), % read alignment (Picard), % of read duplication (Picard). The biological covariates were donor identity, age, sex, brain region and stimulation.

### Differential Expression Analysis

Differential expression analysis was performed between conditions using the R package Differential expression for repeated measures (DREAM) from VariancePartition (Hoffman et al. 2021). DREAM uses a linear mixed model to increase power and decrease false positives for RNA-seq datasets with repeated measurements. The count matrix and covariate file were used as input. Expression data was normalized using the function voomWithDreamWeights, which includes voom transformation. We modeled donor identity as a random effect, since each donor can contribute multiple brain regions, and added selected covariates to adjust for possible technical and biological confounders. The model accounted for sex, donor ID, age, brain region, percentage of mRNA bases, percentage of ribosomal bases, percentage of read duplication, and percentage of reads aligned. *P*-values were then adjusted for multiple testing using the Benjamini-Hochberg False Discovery Rate (FDR) correction.

#### Stimulation

Differential stimulation analysis was performed with DREAM as described above. After this, we used a contrast matrix to test the difference between stimulated sample and cultured sample at baseline. Multiple contrasts were evaluated at the time of the model fit and the results were extracted with FDR < 0.05 for each stimulation (compared to baseline) separately.

#### Age

Differential age analysis was performed with DREAM as described above in all microglia *cultured* samples at the same time. Subsequently, we tested for statistical enrichment using Fisher exact test at FDR < 0.05 comparing the genes associated with aging in the *cultured* microglia samples and the stimulation specific up- and downregulated genes in LPS, R848, IFNγ, DEX, IL-4 and ATP separately. We highlighted aging genes that overlap with MiGA (Lopes et al. 2022).

#### Sex

Differential sex analysis was performed with DREAM as described above in all microglia *cultured s*amples at the same time. Sex analyses were performed on autosomal genes only. We did not perform subsequent analyses because results were not significant.

### Differential Transcript Usage

Transcript expression was estimated in each sample using RSEM with the GENCODE v30 transcript reference. Lowly expressed transcripts were removed with the threshold transcript counts per million > 1 in at least 30% of all samples. Differential transcript usage was tested simultaneously between each stimulation using satuRn, a fast method for computing differential transcript usage. No current differential transcript usage tool can model random effects so we were unable to account for shared donors, but otherwise, the same technical covariates were used as in the differential expression modeling. Pairwise comparisons between baseline and stimulation were extracted using the limma::makeContrasts() function. To correct for test statistic inflation due to correlation across transcripts and donors, we employed a more stringent empirical FDR correction. Transcripts were considered differentially used between stimulation at an empirical FDR < 0.05.

### Pathway analysis

We performed canonical pathway analyses in the Ingenuity Pathway Analysis (IPA) software independently using the following input gene sets: up- and downregulated genes in LPS, IFNγ, R848, IL-4, dexamethasone, ATP at FDR < 0.05 with a log fold change of < -1 or > 1. We also analyzed canonical pathways associated with splicing in the stimulation differential transcript usage (DTU) gene set at empirical FDR < 0.05 using IPA and Gprofiler2 package (Reimand et al. 2016). For the brain region analyses, gene ontology biological process terms were tested for enrichment using the Gprofiler2 package (Reimand et al. 2016). Enriched terms were manually compared between MFG and SVZ, and unique terms were grouped into sets for presentation.

### External Datasets

Raw gene counts files were downloaded for cultured monocytes stimulated with LPS and IFNγ. Differential stimulation analysis was performed with DREAM as described above. We added selected covariates to adjust for possible technical and biological confounders as described before (Navarro et al. 2021). We decided to use a design which includes those covariates that explained the most variance in gene expression which is as follows: *expression* ∼ *rna_batch + age + sex + donor identity + MEDIAN_3PRIME_BIAS + PCT_INTERGENIC_BASES + MDS1 + MDS2 + MDS3 + MDS4*. *P*-values were then adjusted for multiple testing correction using the Benjamini-Hochberg False Discovery Rate (FDR) correction. After this, we used a contrast matrix to test the difference between stimulated sample and *cultured* sample at baseline. Multiple contrasts were evaluated at the time of the model fit and the results were extracted with FDR < 0.05 for each stimulation (compared to baseline) separately.

To compare our LPS findings we also downloaded processed lists of DEGs from a previous microglia study in mice (Gerrits et al. 2020)

### Genotype Quality Control and Imputation

Samples were genotyped using the Illumina Infinium Global Screening Array (GSA) plus a custom disease SNP content (∼60,000 SNPs) for a total of ∼700,000 common variants. To select high-quality data, we applied an initial genotyping quality control, keeping SNPs with call rate > 95%, minor allele frequency (MAF) > 5%, Hardy-Weinberg equilibrium (HWE) P-value > 1 x 10-6, and sample call rate > 95%.

Genotype imputation was performed through the Michigan Imputation Server v1.4.1 (Minimac 4) using the 1000 Genomes (Phase 3) v5 (GRCh37) European panel and Eagle v2.4 phasing in quality control and imputation mode with rsq filter set to 0.3. Following imputation, variants were lifted over to the GRCh38 reference to match the RNA-seq data using Picard liftoverVCF and the “b37ToHg38.over.chain.gz” liftover chain file. Finally, we applied another round of variant quality controls, removing indels and multi-allelic SNPs, and keeping only variants with Hardy-Weinberg P value >1×10^-6^. These variants were additionally annotated using dbSNP (All_20180418.vcf.gz) and snpEff v4.3i. This resulted in 53,502,233 genotyped variants passing all QC steps in 100 donors, of which 4,514,084 had MAF > 0.1.

DNA samples were matched to the RNA-seq data to confirm the same donor origin using the MBV tool from QTLtools and sex mismatching samples were removed by comparing DNA inferred sex from PLINK to RNA gene expression of the *UTY* and *XIST* genes. 16 potential sample swaps were removed. Genetic ancestry of samples was confirmed by principal components analysis using Somalier, overlaying samples with those from the 1000 Genome Project samples (Phase 3). 64 of the 67 donors were confirmed to be of European ancestry, with 3 donors being admixed. Pairwise genetic relatedness was checked using Somalier. No donors were related beyond IBS > 0.125.

### Quantitative Trait Loci mapping

To resolve functional causal variants and genes, we mapped *cis*-eQTLs by testing for association between imputed common genetic variants (MAF > 0.1) and gene expression of a nearby gene in the microglial transcriptome profiles. For the two largest brain region cohorts (MFG and SVZ) we first mapped eQTLs in the unstimulated, LPS and IFNγ conditions separately. For each QTL mapping round, we included 3 PEER factors to account for technical variation in gene expression and the top 5 principal components of the genotype matrix from Somalier. For each gene, we included all SNPs at MAF > 0.1 with a 1 megabase window around the transcription start site. Mapping was performed using tensorQTL (https://github.com/broadinstitute/tensorqtl). Multiple testing was accounted for by using a permutation testing scheme in tensorQTL for each gene followed by Storey’s q-value correction of the top permutation P-value for each gene. Due to the small size of the cohorts, the false discovery rate was set at a permissive 20%.

We then used the Suez framework to map response cis-eQTLs (https://github.com/davidaknowles/suez). Suez is an extension of the PANAMA (Fusi, Stegle, and Lawrence 2012) linear mixed model (LMM) framework to map eQTLs and response eQTLs while accounting for latent confounding. A detailed explanation of the methods can be found in the publication (Knowles et al. 2018). Suez mapping included both baseline and stimulated samples and accounted for mixed ancestries and repeated donors by modeling a genotype relationship matrix as a random effect in the linear mixed model. Response eQTLs were measured by the interaction term between genotype and stimulation. We applied the same thresholds for MAF (10%) (1Mbp) and cis-windows and included 3 principal components of gene expression after comparing the reQTL discovery rate from different numbers of PCs (**Supplemental Figure 13)**.

### Visualization

All plots were created using ggplot2 in R (version 3.6.0), with ggrepel, ggfortify and ggbio or additional layers of visualization.

## Author contributions

LDW, JH, TR conceived and supervised the study. GJLS, MAMS isolated microglia at University Medical Center Utrecht, the Netherlands. GJLS, FAG, RM, and RK isolated microglia at Mount Sinai School of Medicine, New York City. GJLS, RK performed genotyping and RNA-seq. KPL performed data pre-processing and quality control. GJLJ led analyses, with input from JH, LW, and TR. JH led genetic analyses. BM assisted with genetic analyses. RAV assisted with QTL mapping and performed genotyping QC. The manuscript was written by GJLS, JH, LDW, and TR, with input from all co-authors. All authors read and approved the manuscript.

## Acknowledgements

This study is financially supported by the nos. NIA R21-AG063130 grant from the NIH. TR is also supported by other grants from the NIH (nos. NIA R01-AG054005, NIA U01-AG068880, NIA RF1-AG065926, NIA R56-AG055824 and NINDS R01-NS116006). Gijsje Snijders was supported through the foundation BBRF in the USA. The authors thank the teams of the Netherlands Brain Bank and the Mount Sinai Neuropathology Brain Bank and Research CoRE for their services. We thank the study participants for their generous gifts of brain donation. The microglia were isolated through the efforts of a large team and we would like to thank Manja Litjens, Roland D. van Dijk, Alba Fernández-Andreu, Paul R. Ormel, Hans C. van Mierlo, Y. He, Stephanie Gumbs, Miriam E van Strien, Saskia Burm, Vanessa Donega, and Elly M. Hol for all their contributions to this effort. We thank the members of the Raj and de Witte labs for their feedback on the manuscript.

## Data Availability

Raw and processed RNA-seq and genotype data sets are deposited in the National Institute on Aging Genetics of Alzheimer’s Disease Data Storage Site (NIAGADS) at https://dss.niagads.org/datasets/ng00105/ Accession number: NG00105.v1. The user will need to log into NIAGADS Data Access Request (DAR) to start an application.

Instructions to download the dataset can be found at https://www.niagads.org/data/request/data-request-instructions.

All differential expression results and gene lists are present as supplementary tables.

## Code Availability

All code used to make figures: https://github.com/RajLabMSSM/MiGASti

Response QTL mapping pipeline: https://github.com/RajLabMSSM/suez-snakemake-pipeline

---

**Supplementary Figure 1.**
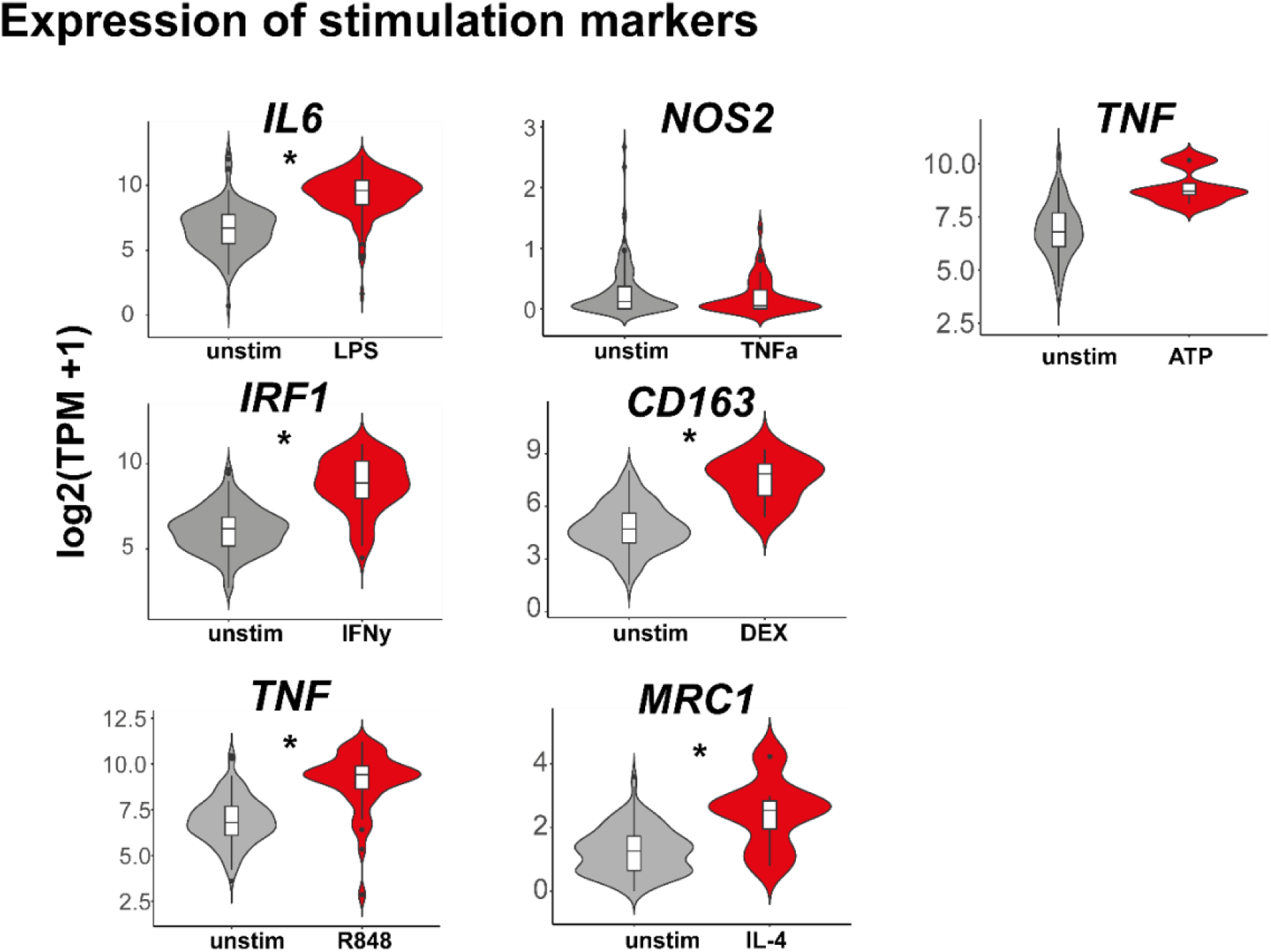
Gene expression of well-known stimulation specific markers across different stimulations. A) Violin plots depicting expression in transcripts per million (TPM) for representative (anti)-inflammatory genes *IL6, IRF1, TNF, NOS2, CD163, MRC1, TNF* that are known to be upregulated after stimulation with respectively LPS, IFNγ, R848, TNF-α, DEX, IL-4 or ATP in the stimulated compared to the unstimulated samples. * Statistical significance based on adjusted P-values (FDR) derived from the differential expression results.

**Supplementary Figure 2.**
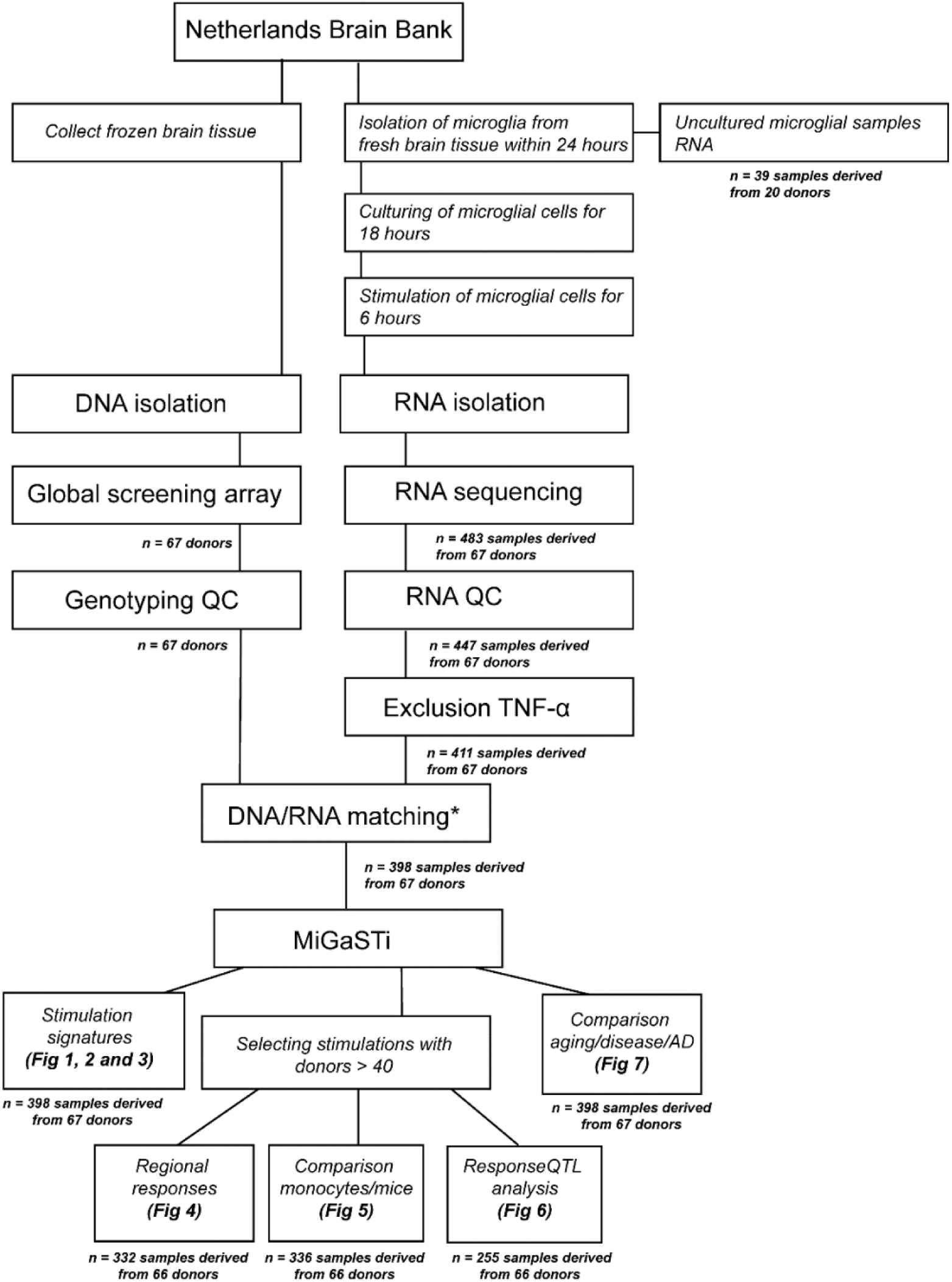
Flowchart of quality control of the microglial samples. Abbrevations QC = quality control. * DNA-RNA sample matching using QTL tools-mbv measured by the percentage of concordance at heterozygous and homozygous genotypes.

**Supplementary Figure 3.**
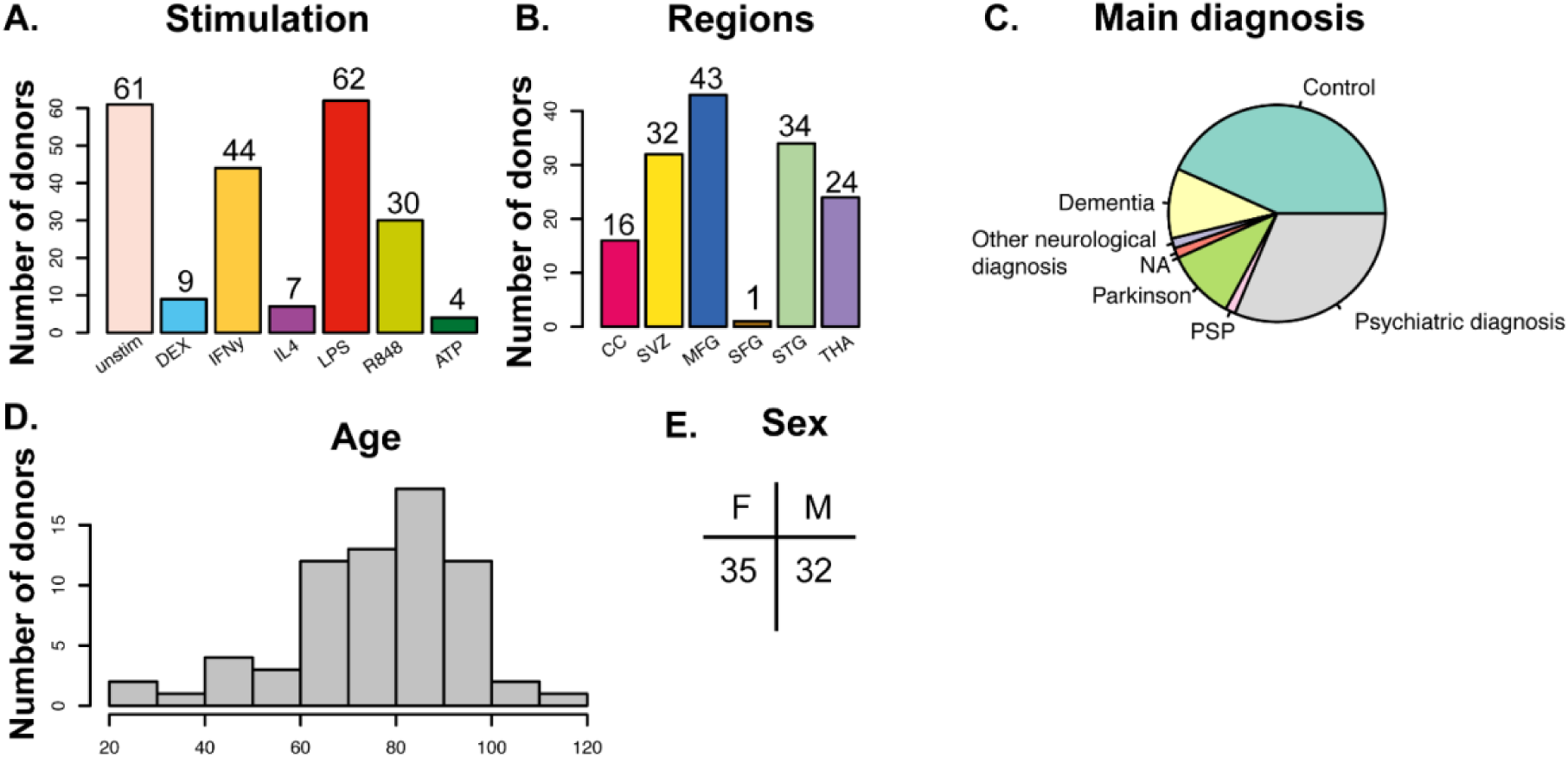
Overview of the data. A) Number of donors by stimulation. B) Numbers of donors by brain region. Each donor donated one up to five brain regions. C) Frequency of diagnosis. D) Age range of the 66 donors in this study. E) The numbers of female (F) and male (M) donors included. Abbrevations ATP: adenine triphosphate, CC: corpus callosum, DEX: dexamethasone, F: female, IFNγ: Interferon gamma, IL4: interleukin-4, LPS: lipopolysaccharide, M: male, MFG: medial frontal gyrus, NA: not applicable, PSP: progressive supranuclear palsy, R848: resiquimod, SFG: superior frontal gyrus, STG: superior temporal gyrus, SVZ: subventricular zone, THA: thalamus, unstim: unstimulated.

**Supplementary Figure 4.**
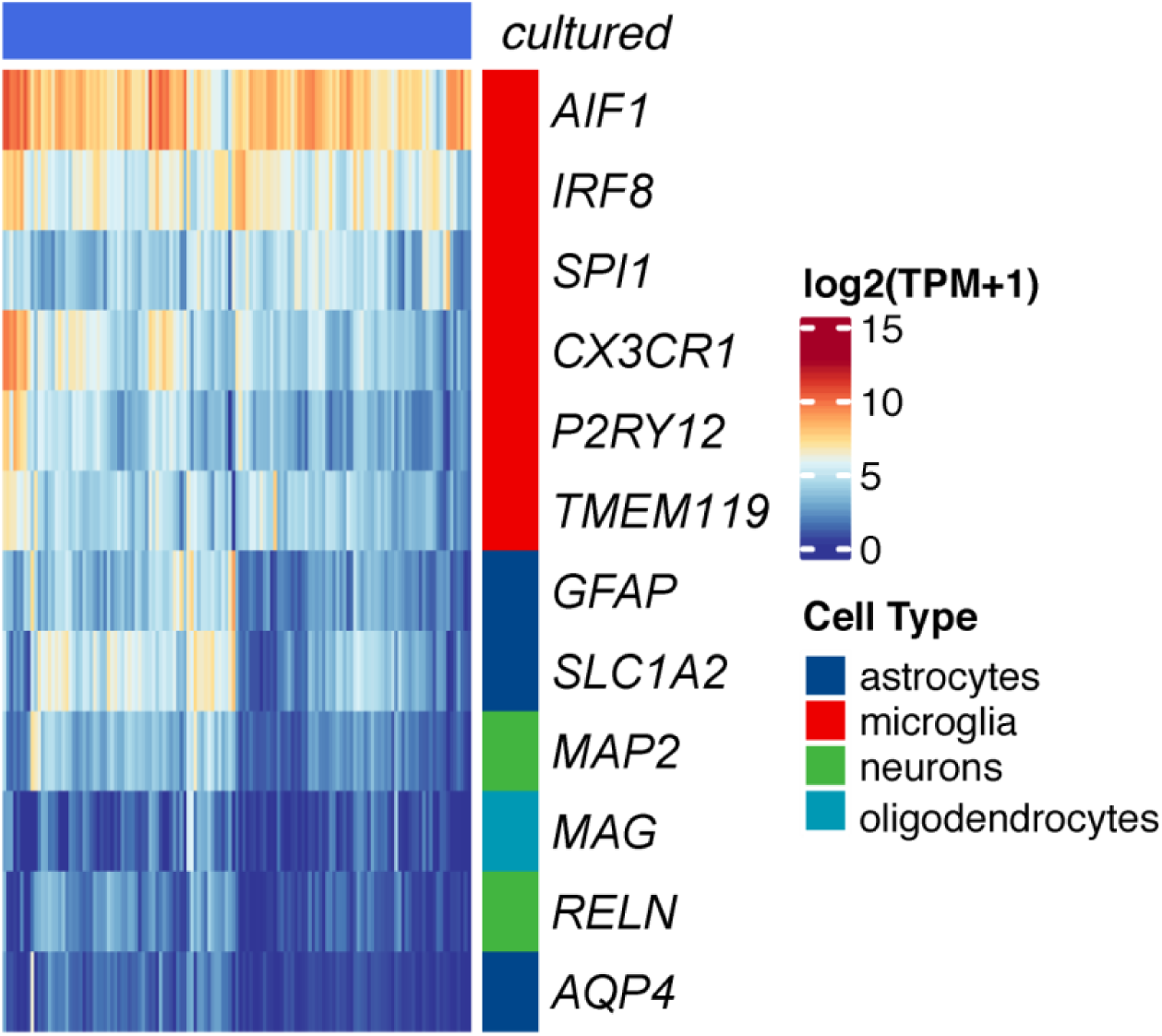
Purity of cultured microglia samples. Expression levels (TPM+1 log2 scale) of cell markers in 398 *cultured* microglial samples. Blue colors indicate low expression and red colors indicate high gene expression.

**Supplementary Figure 5.**
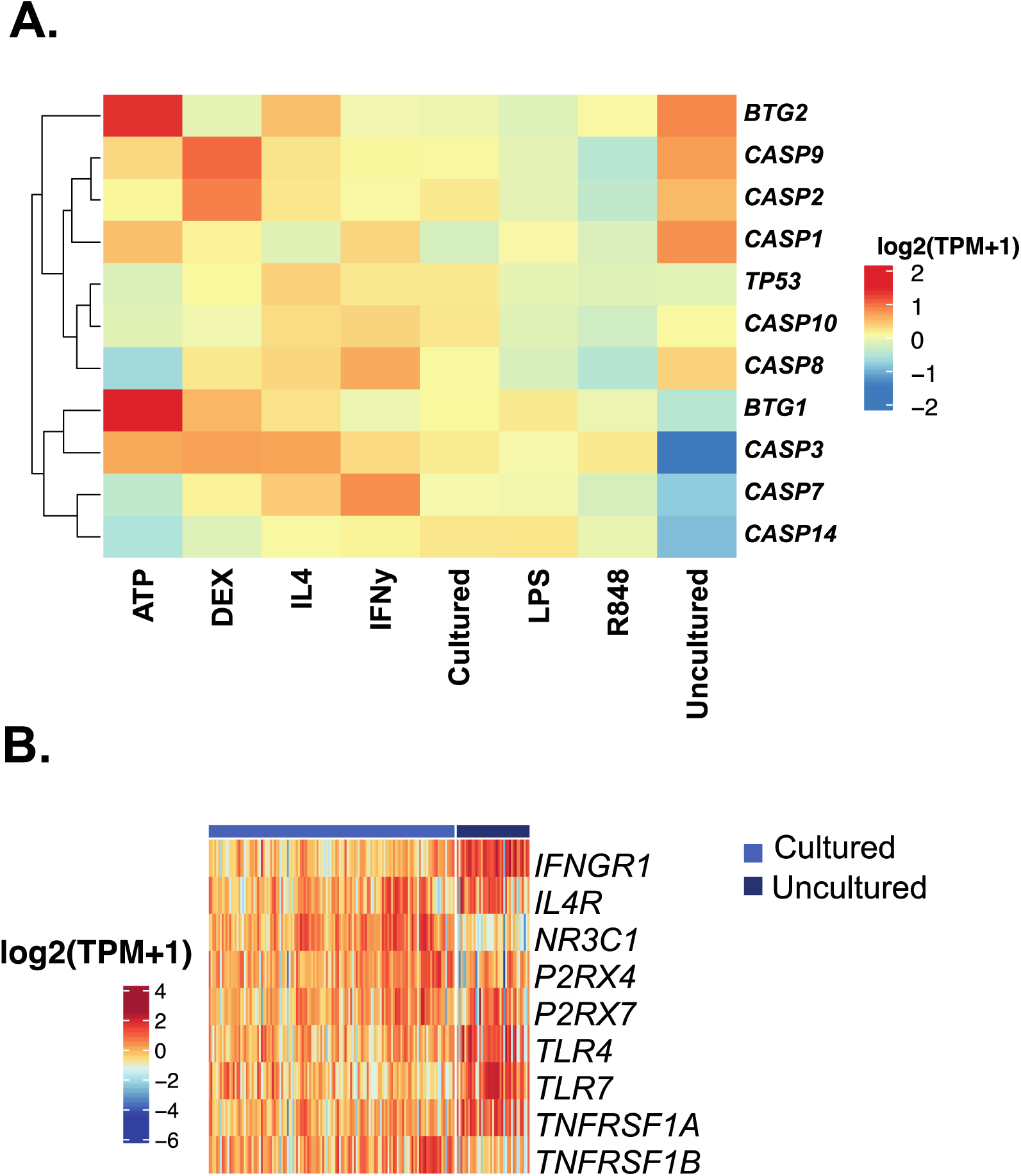
Expression of known apoptotic markers and receptors for inflammatory triggers. A) Heatmap depicting the average expression (TPM+1 log2 scale) of caspases and apoptotic cell markers for each stimulation separately compared to the cultured/uncultured condition. Clustering dendrogram is based on Euclidean distances. B) Expression levels (TPM+1 log2 scale) of known receptor markers for inflammatory triggers for cultured versus uncultured samples. Blue colors indicate low expression and red colors indicate high gene expression.

**Supplementary Figure 6.**
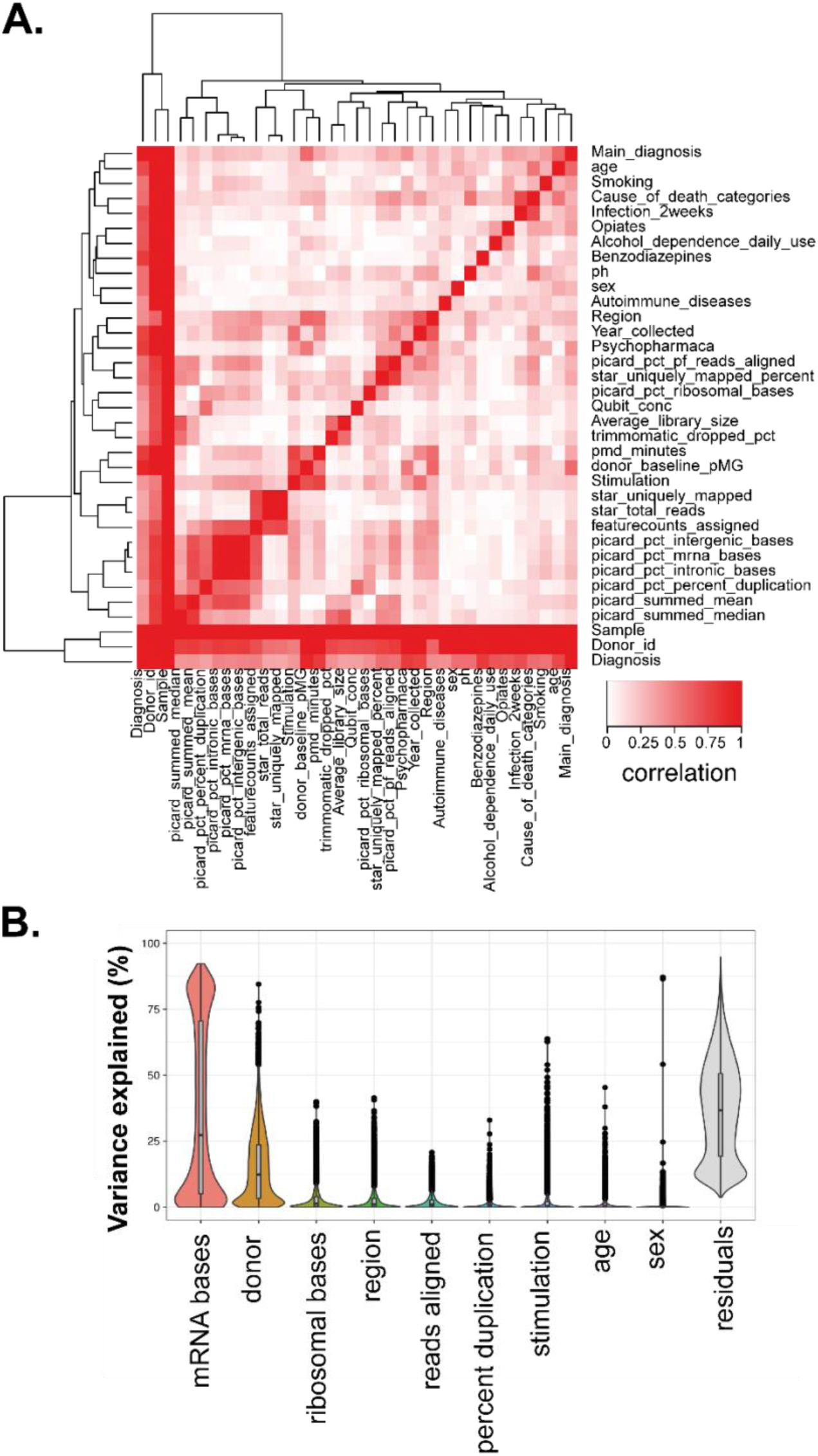
Main sources of expression variation and correlation of covariates. A) Pairwise canonical correlation between the covariates. Red indicates high correlation; white indicates low correlation. B) VariancePartition computes variance explained per gene for each variable independently. Boxplots show the median, first and third quartile of the distribution. Jittered points are genes outside 1.5 times the interquartile range. More variance is explained on average by technical factors than clinical factors.

**Supplementary Figure 7.**
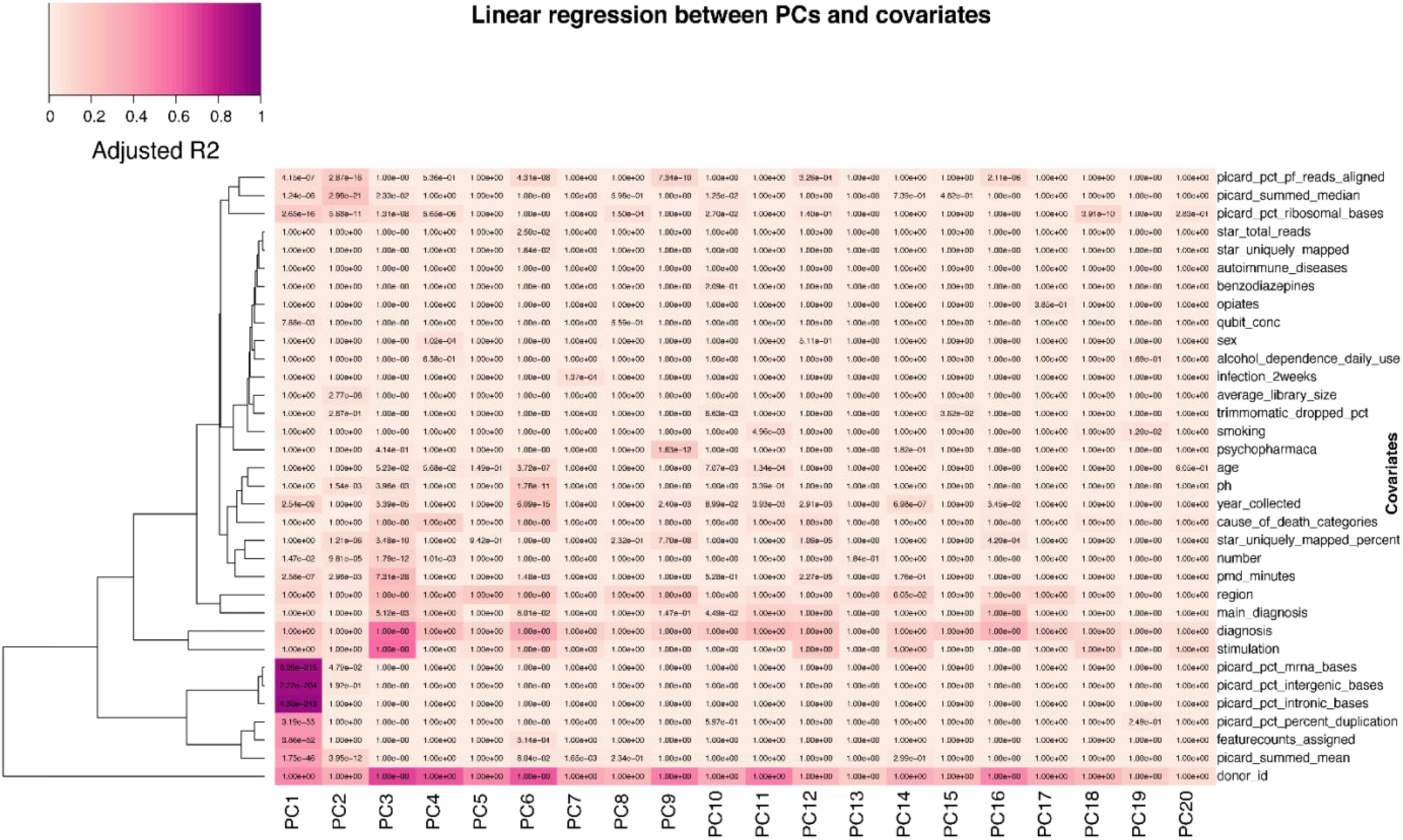
Sources of variation in the gene expression data. Linear regression between the first 20 Principal Components (PCs) and the covariates. Colors correspond to the adjusted R-squared. The P-values are from the Linear regression, two-sided and Bonferroni adjusted.

**Supplementary Figure 8.**
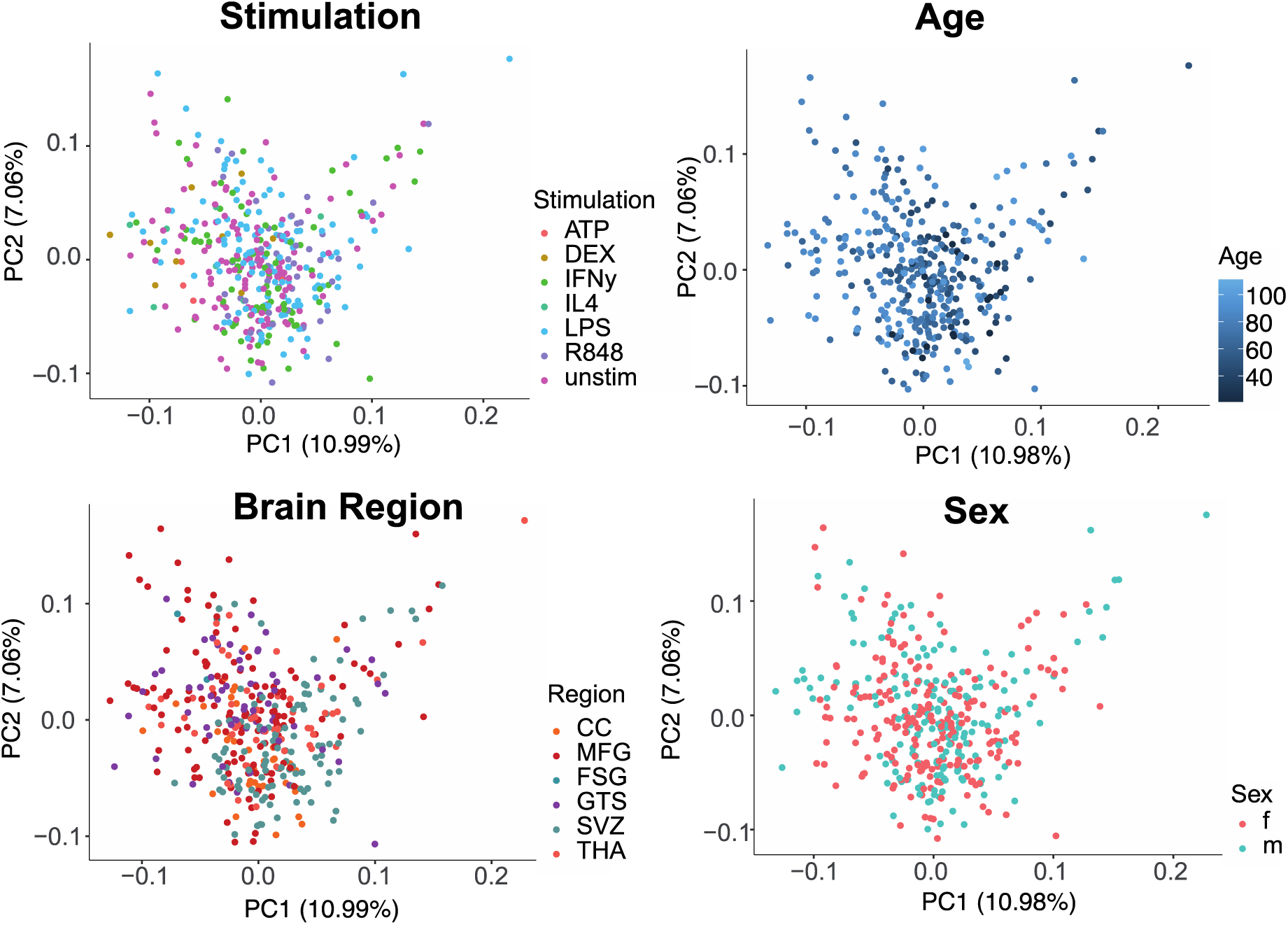
Principal component analysis (PCAs) after data adjustment. PCA after correction by regressing out technical confounders, colored by stimulation, age, brain region and sex. The plots show voom-TMM normalized expression for all 398 cultured microglia samples.

**Supplementary Figure 9.**
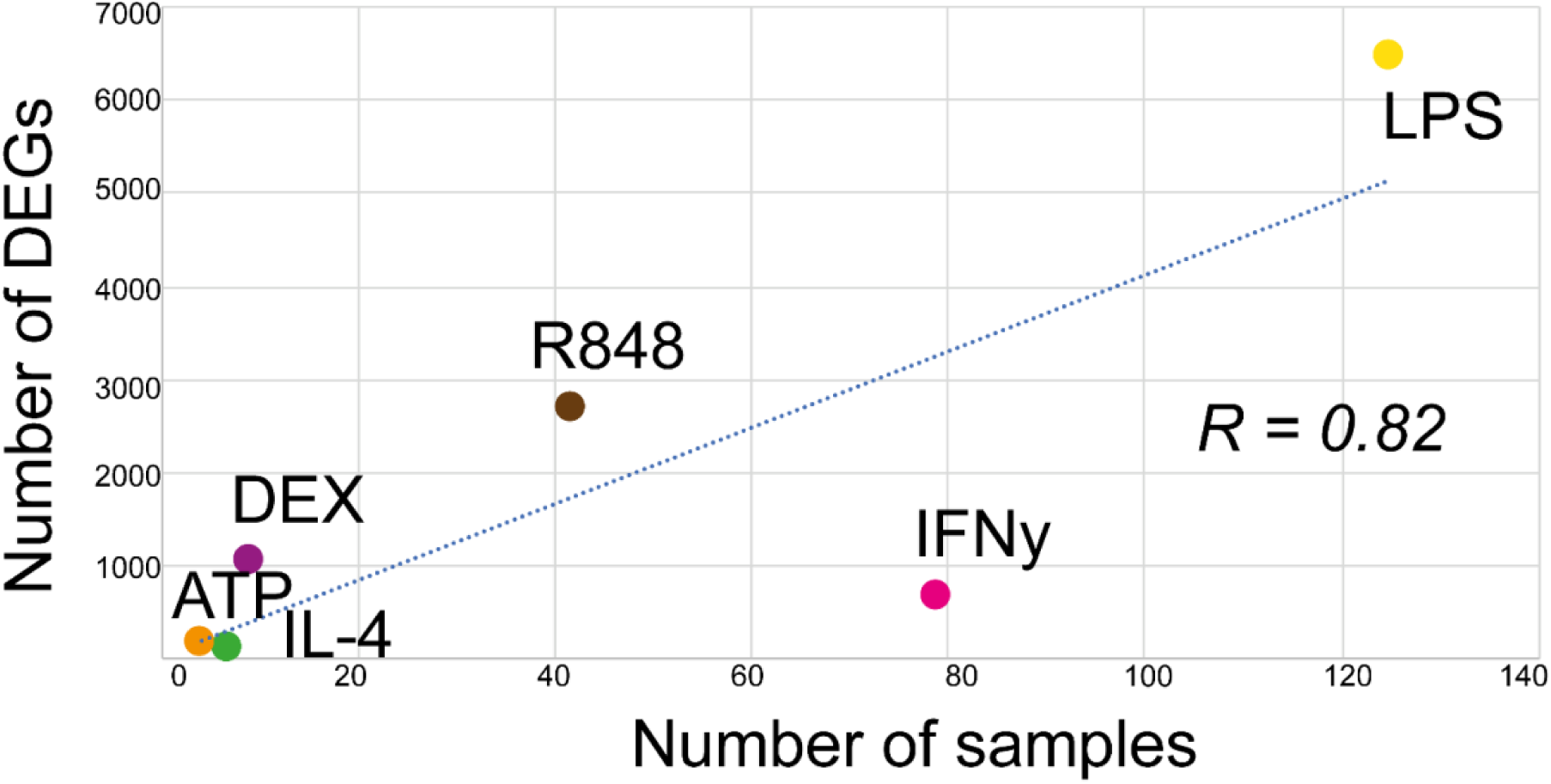
Correlation number of samples and number of differentially expressed genes (DEGs). The linear correlation is calculated with Pearson correlation coefficient (R). Abbrevations. ATP: adenine triphosphate, DEX: dexamethasone, IFNγ: Interferon gamma, IL4: interleukin-4, LPS: lipopolysaccharide, R848: resiquimod.

**Supplementary Figure 10.**
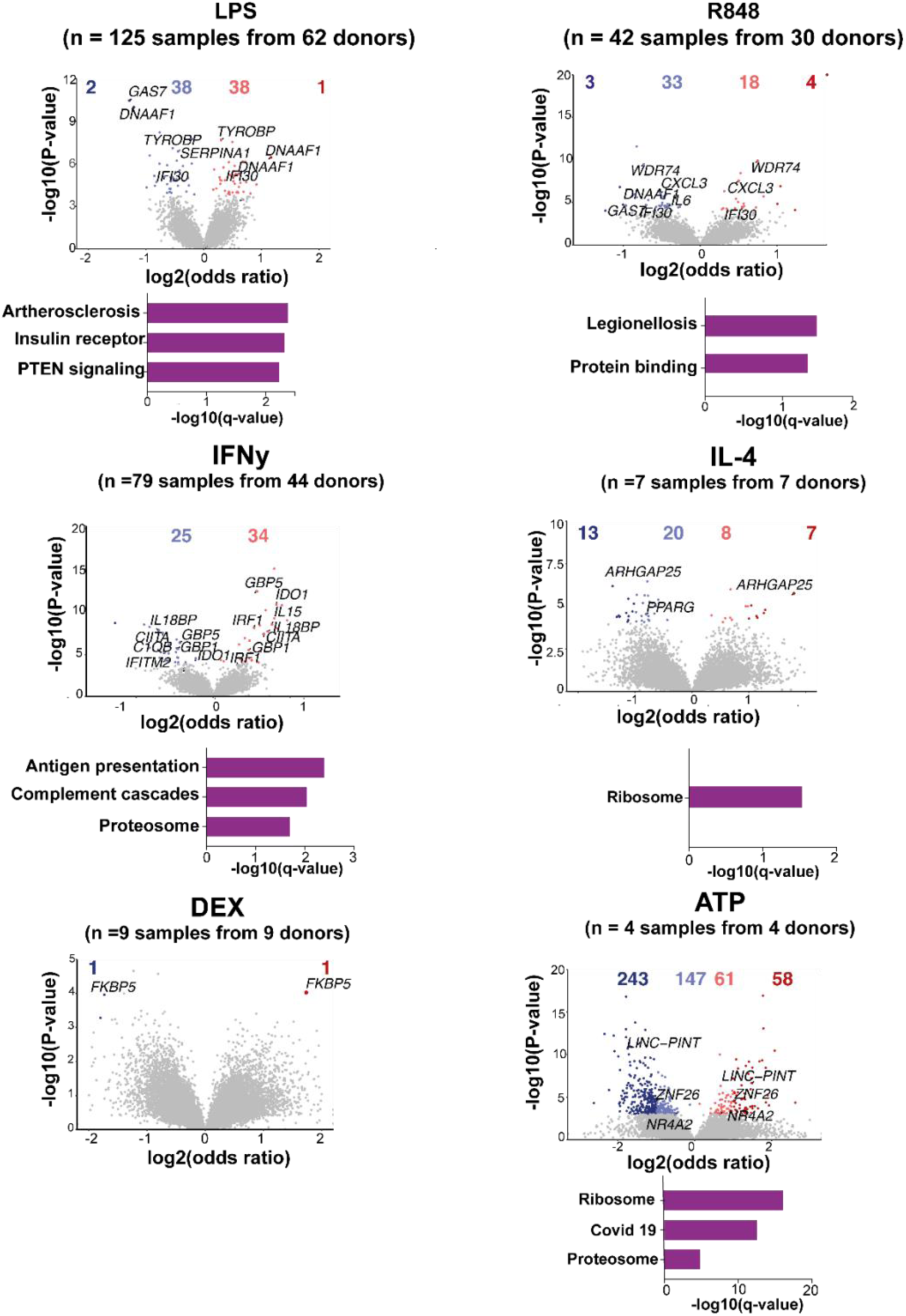
Differential transcript usage (DTU) across stimulations. Volcano plots comparing stimulated microglia to controls in all donors. Genes colored by whether or not differentially expressed (empirical FDR < 0.05; grey), differentially expressed but with modest effects (| odds ratio | 1; orange/blue) and with stronger effects (| odds ratio | > 1; red/dark blue). Numbers of genes in each category above the plot. Genes related to Alzheimer’s disease or stimulation response genes are highlighted. Functional enrichment analyses of all differential expressed genes using IPA or gProfiler2. Significantly enriched terms are shown with q-value 0.05.

**Supplementary Figure 11.**
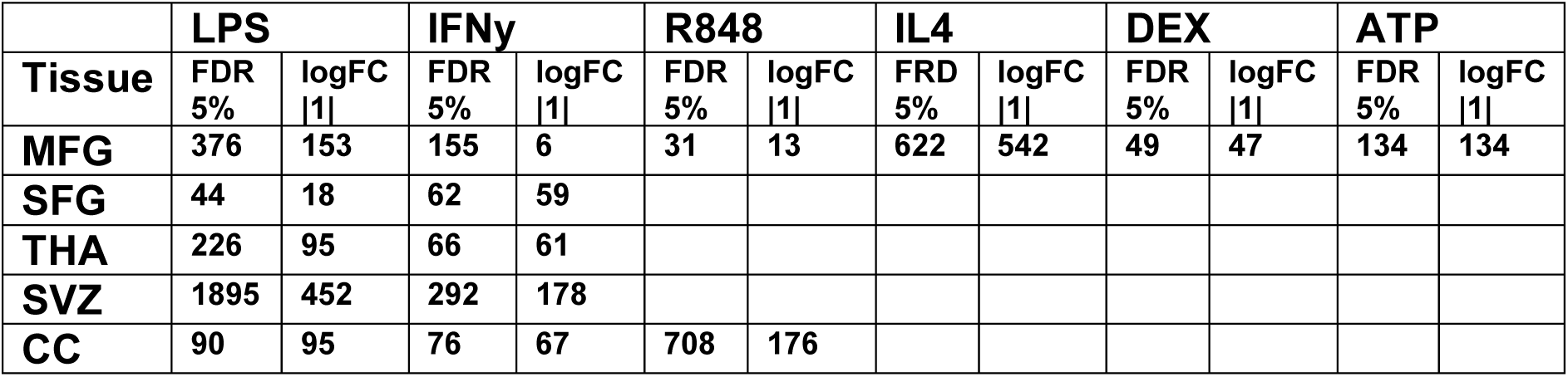
Number of differentially expressed genes across brain regions in different stimulations. Abbrevations ATP: adenine triphosphate, CC: corpus callosum, DEX: dexamethasone, FDR: false discovery rate, IFNγ : Interferon-gamma, IL4: interleukin-4, logFC: log fold change, LPS: lipopolysaccharide, MFG: medial frontal gyrus, R848: resiquimod, STG: superior temporal gyrus, SVZ: subventricular zone, THA: thalamus,

**Supplementary Figure 12.**
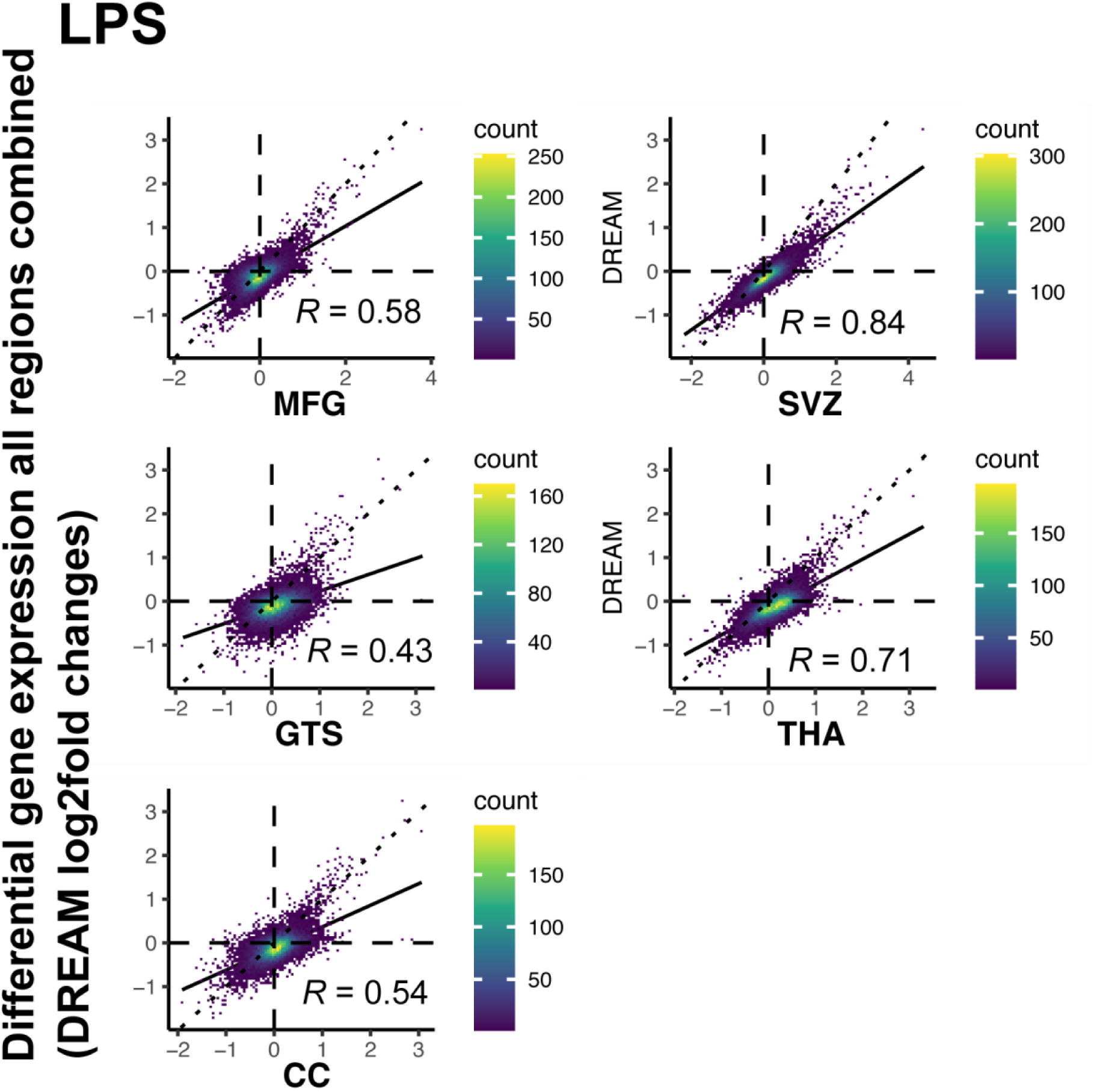
Correlating differential gene expression between each brain region separately and all brain regions combined in LPS. Scatter plots showing pairwise correlations in LPS between differential gene expression results of each brain region separately and all brain regions combined of log_2_ fold changes using Pearson correlation. Individual points are genes, color refers to density of overlapping points.

**Supplementary Figure 13.**
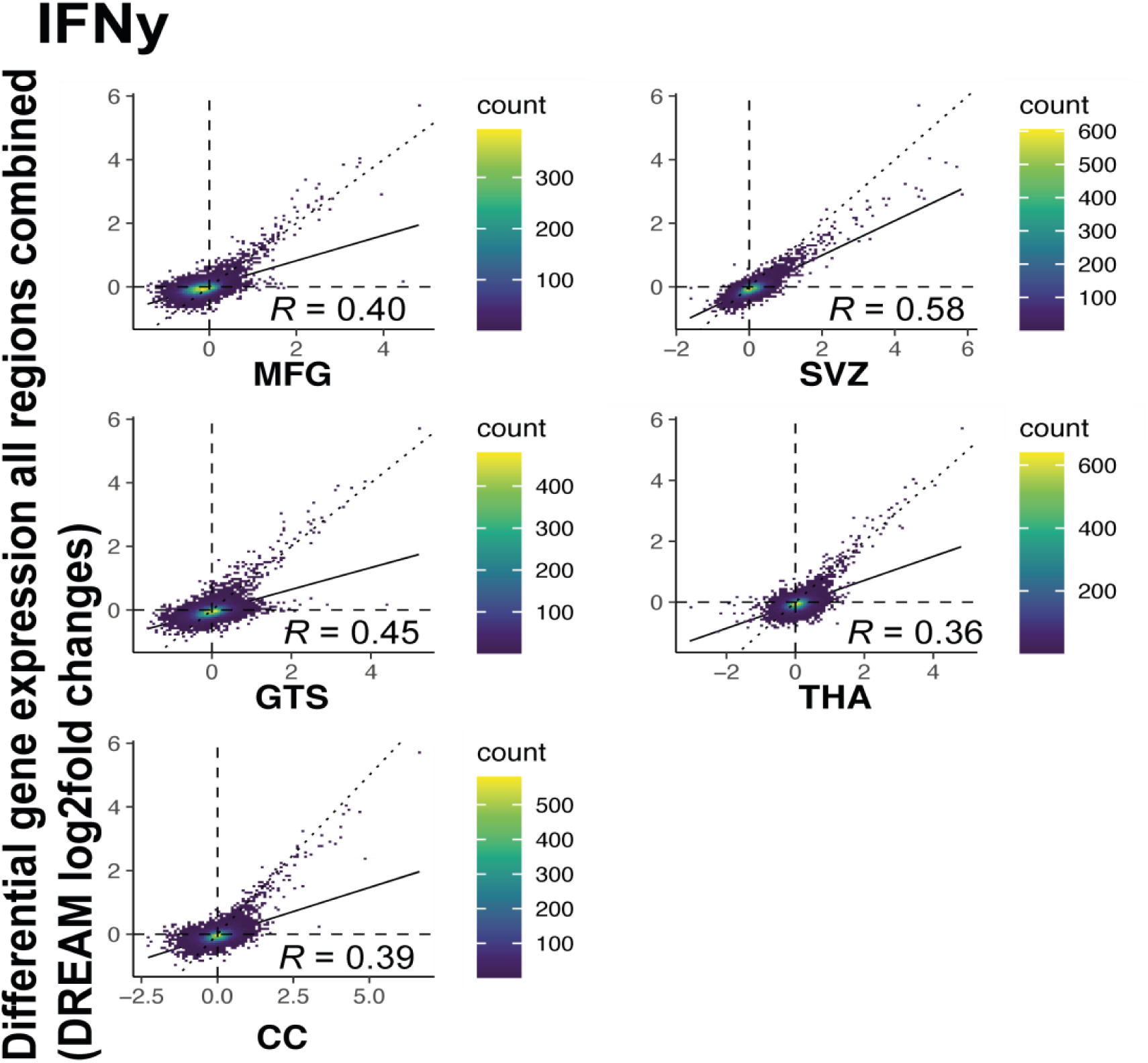
Correlating differential gene expression between each brain region separately and all brain regions combined in IFNγ. Scatter plots showing pairwise correlations in IFNγ between differential gene expression results of each brain region separately and all brain regions combined of log_2_ fold changes using Pearson correlation. Individual points are genes, color refers to density of overlapping points.

**Supplementary Figure 14.**
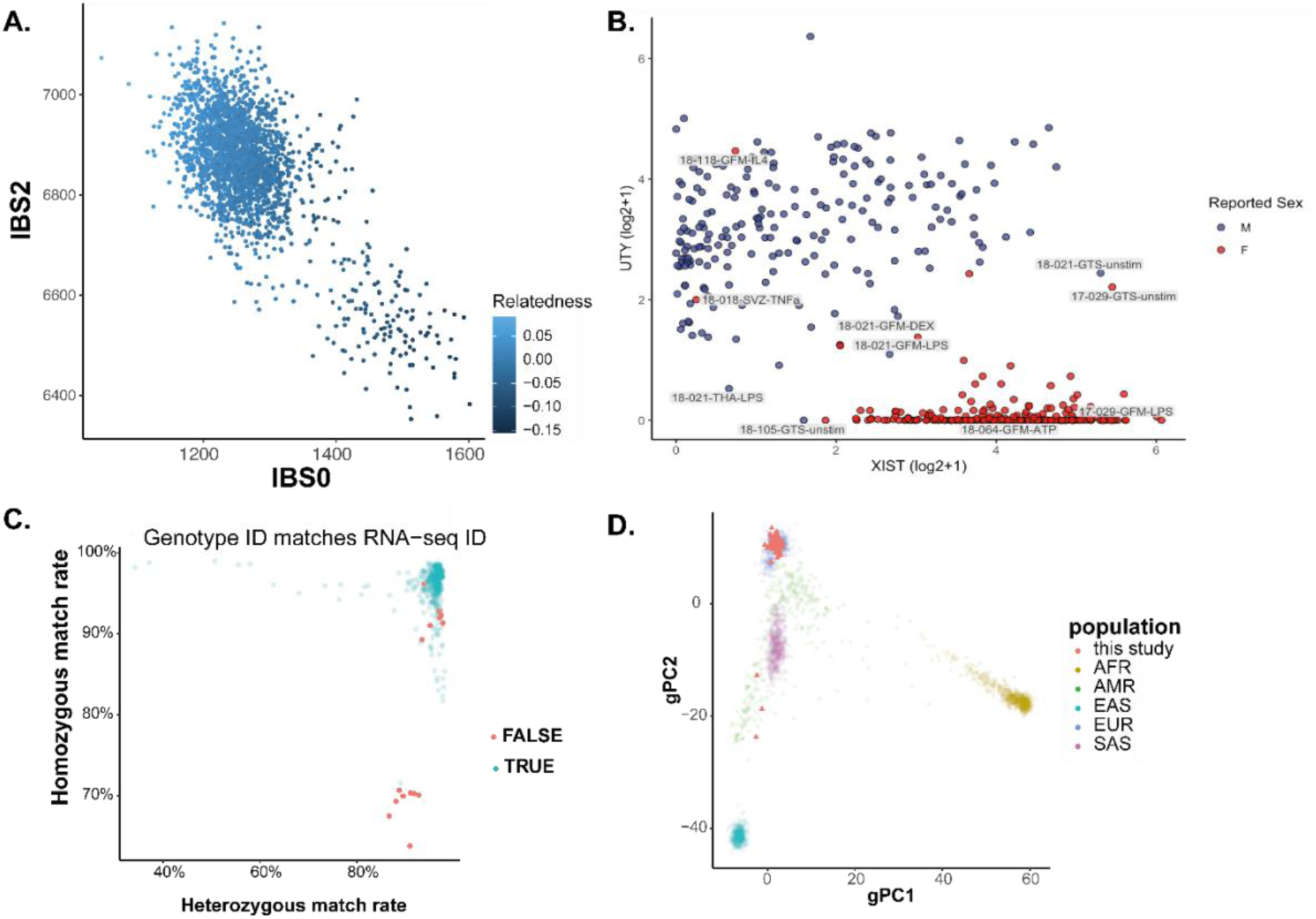
Genotyping QC. A) Relatedness estimates from Somalier. B) Log scaled expression (voom) of sex chromosome genes (UTY and XIST) for each donor colored by reported sex (blue = male, red = female) C) DNA-RNA sample matching using QTLtools-mbv measured by the percentage of concordance at heterozygous and homozygous genotypes. Samples in red failed to match ids between each data. D) First two ancestry principal components of MiGASTi samples (red squares) on top of 1000 Genome individuals colored by major populations. African [AFR], Admixed American [AMR], East Asian [EAS], European [EUR], South Asian [SAS]

**Supplementary Figure 15.**
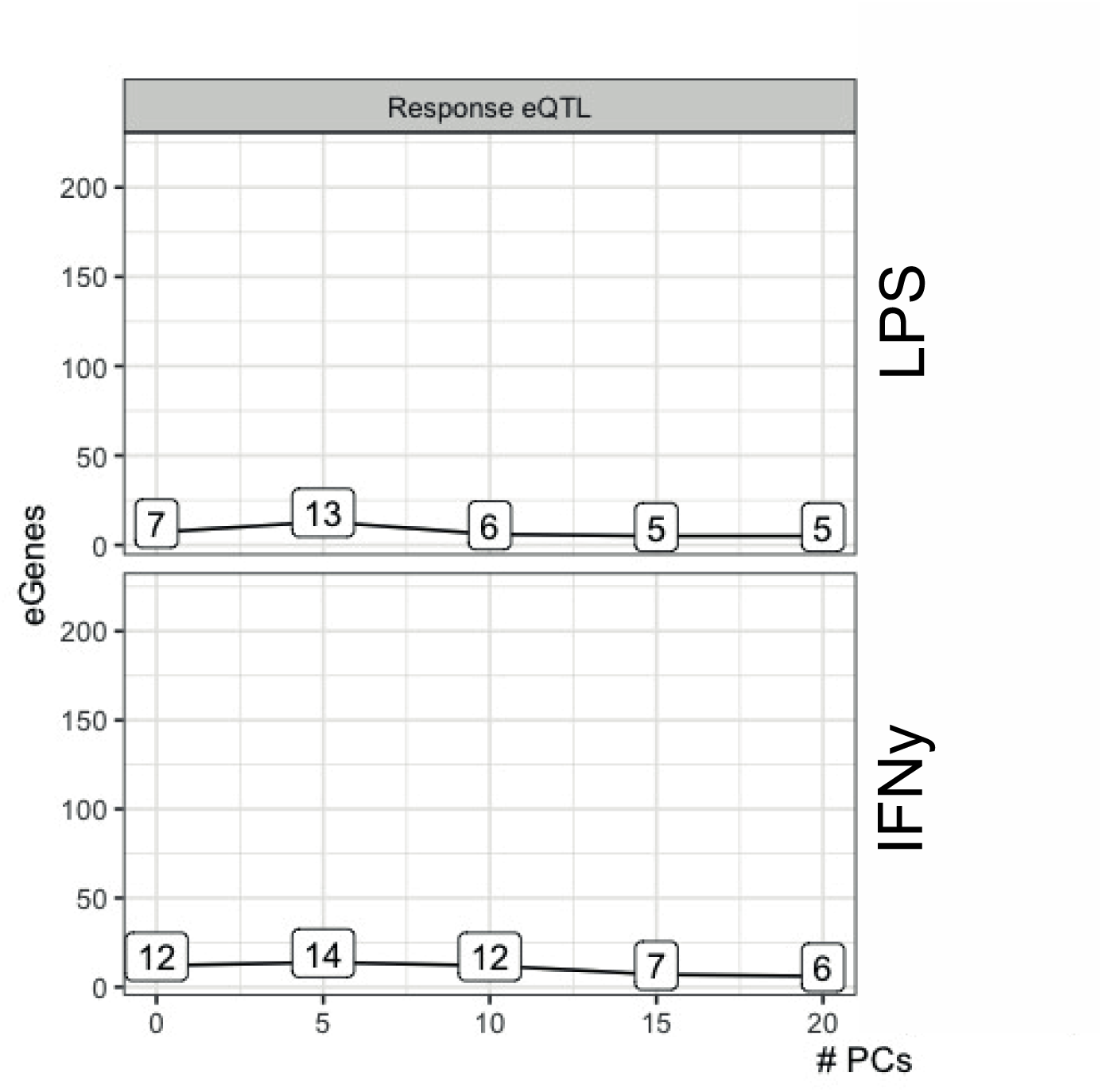
responseQTL discovery rate with different numbers of principal components. Upper panel - number of genes with an reQTL at qvalue < 0.05 in LPS and PC threshold. Lower panel - number of genes with an reQTL at qvalue < 0.05 in IFNg and PCthreshold.

**Supplementary Figure 16.**
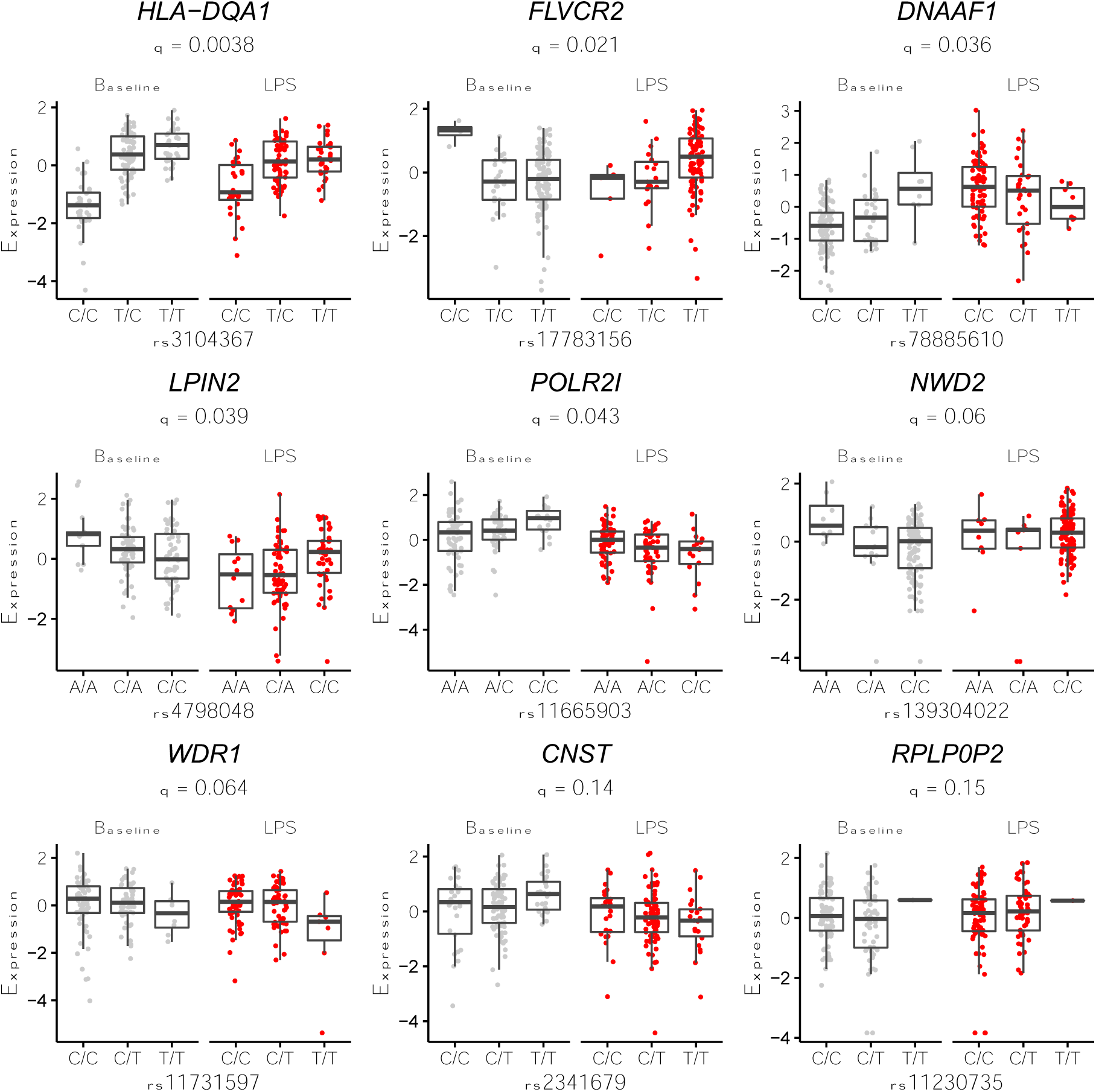
Mapping response cis-eQTLs (reQTLs) in microglia stimulated with LPS. Boxplots show interaction between genotype and stimulation on residualized gene expression levels. q refers to adjusted P-value following correction for numbers of tested variants and genes.

**Supplementary Figure 17.**
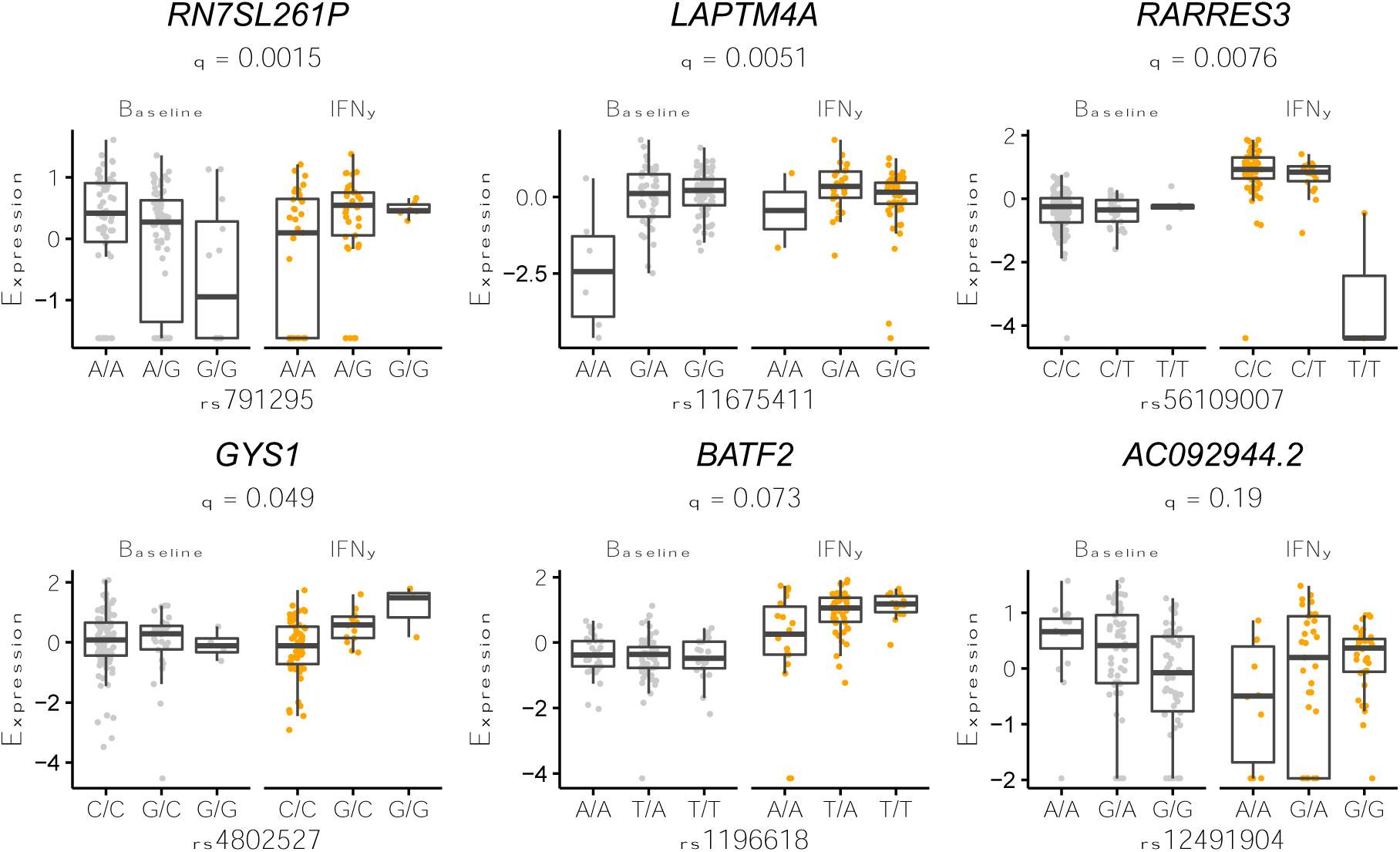
Mapping response cis-eQTLs (reQTLs) in microglia stimulated with IFN-γ. Boxplots show interaction between genotype and stimulation on residualised gene expression levels. q refers to adjusted P-value following correction for numbers of tested variants and genes.

**Supplementary Figure 18.**
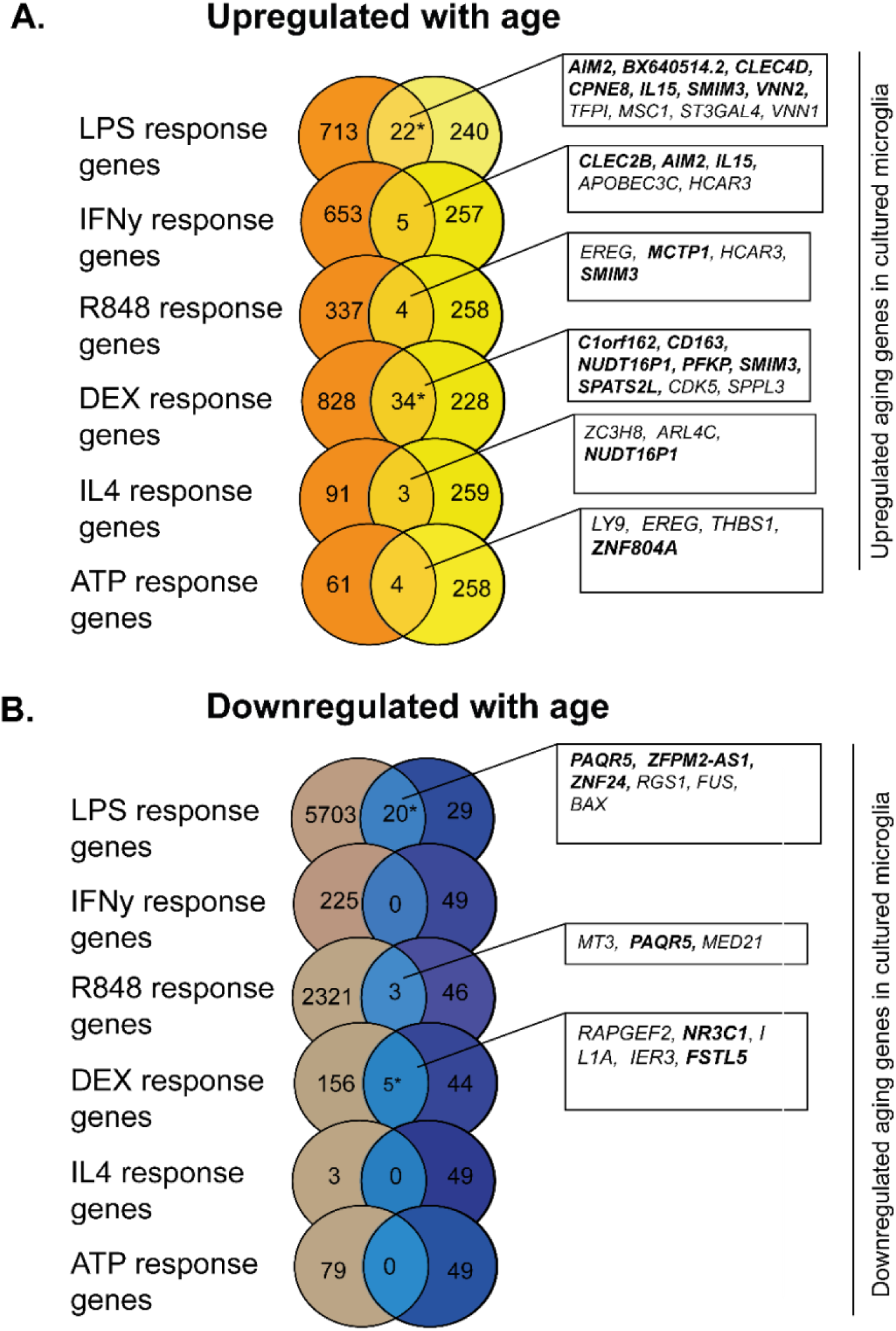
Overlap of age-related genes and stimulation response genes in MiGASTi. Enrichment analysis between differentially expressed aging genes with stimulation response genes in MiGASTi. A) *Upregulated in age between LPS and DEX in MiGASTi. Asterisk indicates significant enrichment by a one-sided Fisher’ exact test (upregulated LPS: OR 2.4, p < 0.0001), upregulated DEX: OR 3.4, p < 0.0001) Genes highlighted in bold were replicated upregulated aging genes in MiGA (Lopes et al. 2022) B) Downregulated in age between LPS and DEX. Asterisk indicates significant enrichment by a one-sided Fisher’ exact test (downregulated LPS: OR 1.7, p = 0.04, downregulated DEX: OR 14.4, p < 0.0001). Genes highlighted in bold were replicated aging genes in MiGA (Lopes et al. 2022).

**Supplementary Figure 19.**
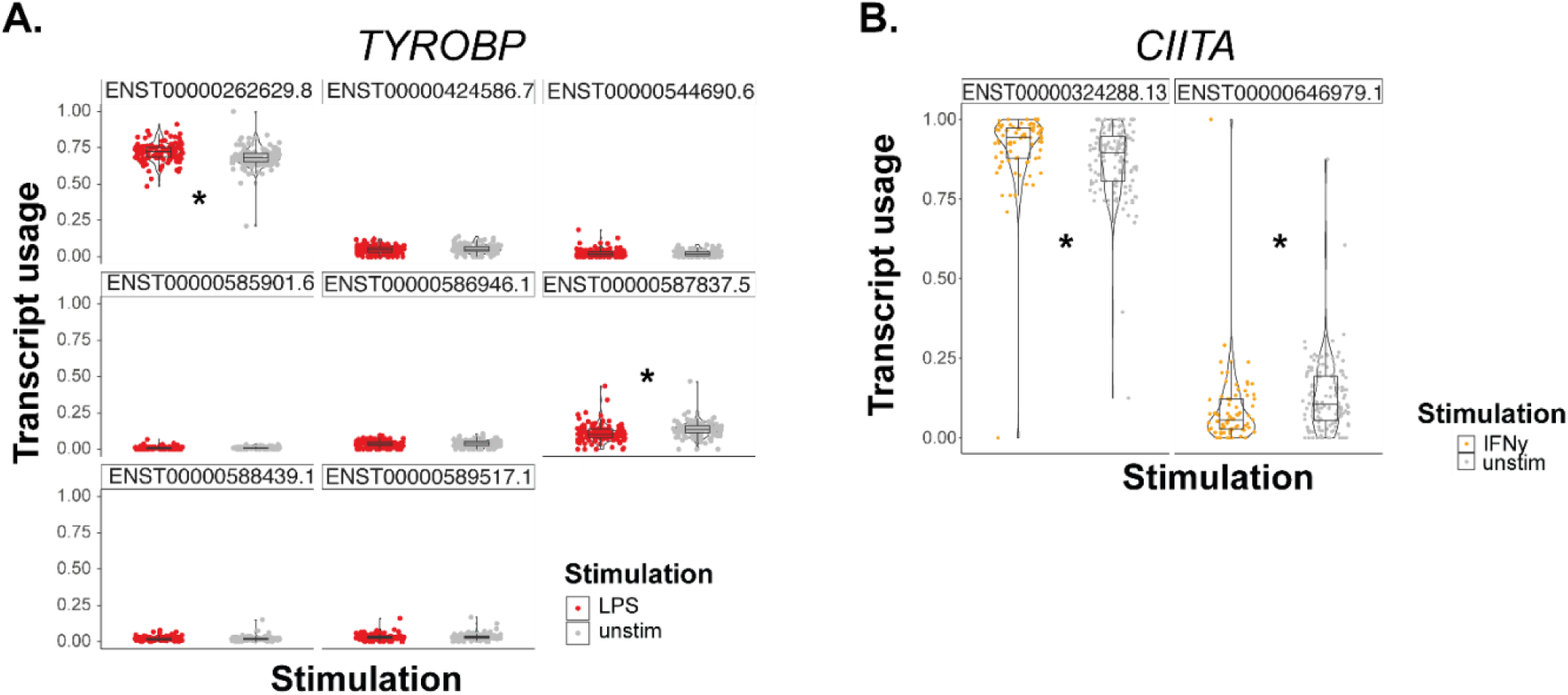
Differential transcript usage in LPS and IFNγ. A) Example transcript usage of *TYROBP* in LPS. Asterisks indicate significance. The DTU signal is driven by a decrease respectively increase in the transcripts ENST00000292629.8 and ENST0000057837.5. B) Example transcript usage of *CIITA* in IFNγ. Asterisks indicate significance. The DTU signal is driven by a significant decrease respectively increase in the transcripts ENST00000324288.13 and ENST00000646979.1.

